# Nucleocapsid 203 mutations enhance SARS-CoV-2 immune evasion

**DOI:** 10.1101/2021.12.20.473471

**Authors:** Xiaoyuan Lin, Weiwei Xue, Yueping Zhang, Beibei Fu, Jakob Trimpert, Na Xing, Dusan Kunec, Wanyan Tang, Yang Xiao, Kaiwen Meng, Shuobo Shi, Haibo Wu, Geng Meng, Zhenglin Zhu

## Abstract

Previous work indicated that the nucleocapsid 203 mutation increase the virulence and transmission of the SARS-CoV-2 Alpha variant. However, Delta later outcompeted Alpha and other lineages, promoting a new wave of infections. Delta also possesses a nucleocapsid 203 mutation, R203M. Large-scale epidemiological analyses suggest a synergistic effect of the 203 mutation and the spike L452R mutation, associated with Delta expansion. Viral competition experiments demonstrate the synergistic effect in fitness and infectivity. More importantly, we found that the combination of R203M and L452R brings in a 3.2-fold decrease in neutralizing titers to the neutralizing serum relative to L452R-only virus. R203M/L452R show an increased fitness after the initiation of global vaccination programmes, possibly associated with the enhanced immune evasion. Another rapidly emerging variant Omicron also bears the 203 mutation. Thus, we proposed that nucleocapsid mutations play an essential role for the rise and predominance of variants in concern.

## INTRODUCTION

It has been two years since the global outbreak of severe acute respiratory syndrome coronavirus 2 (SARS-CoV-2) in 2019 ^1-5^. By December 2021, SARS-CoV-2 had caused more than 263 million confirmed infections and more than 5 million deaths worldwide. SARS-CoV-2 is an RNA virus with a moderate mutation rate ^4,6,7^, however, the large number of human infections fueled a rapid evolution of this virus. The resulting genomic variations enabled selection of several distinct variants of concern (VOCs), including Alpha (B.1.1.7), Beta (B.1.351), Gamma (P.1/B.1.1.28), Kappa (B.1.617.1), Delta (B.1.617.2) and the newly emerging Omicron (B.1.1.529).

SARS-CoV-2 is an enveloped, positive-stranded RNA virus. Receptor recognition and fusion processes are mediated by the trimeric spike (S) glycoprotein. The S protein is the major target for the development of vaccines and neutralizing antibodies ^8,9^. Adaptive mutations, such as D614G ^10-14^, N501Y ^15-18^, E484K ^19,20^ and L452R ^21,22^, accumulating in the S protein among different VOCs are regarded as essential to increase infectivity, transmissibility and resistance to neutralization.

In addition to these adaptive S protein mutations, there are adaptive mutations in other parts of the viral genome that contribute to the virulence, spread and immune evasion of the virus. Several researchers have highlighted the potential importance of mutations in the nucleocapsid (N) protein ^12,23-25^. Our previous work has found that the cooccurring mutations R203K/G204R in the N protein increase fitness and infectivity of SARS-CoV-2 and facilitate the emergence of the Alpha variant ^23^. The R203K/G204R mutations, dominant in the worldwide pandemic until May 2021, enhance the fitness of SARS-CoV-2 and are associated with the increased transmission and virulence of the Alpha variant. During the global vaccination campaign, Delta has outcompeted other lineages and became predominant worldwide. The Delta variant was first identified as a VOC in India in 2020 ^26^, bearing the S protein mutation L452R, which confers escape from cellular immunity ^21,26^. However, the N protein mutation R203M is likewise associated with the emergence of Delta variants. Two independently evolved VOCs (Alpha and Delta) have mutations located at the same position of the N protein, suggesting functional importance of the N protein 203 mutation for viral adaptation and fitness. Thorough research on the evolutionary and functional effects of R203M is important for understanding the effects of N protein mutations and their contribution to the rapid global rise of the Delta variant.

For these reasons, we evaluated the evolution of the R203M mutation based on all documented SARS-CoV-2 genomic sequences. The results show that this mutation has rapidly expanded since January 2021 with the emergence of Delta (Fig. S1A, B) and is becoming dominant in the worldwide pandemic (Fig. 1A, B). Moreover, our results suggest that the R203M N protein mutation and the L452R S protein mutation have a statistically significant synergistic effect towards the emergence and spread of the Delta variant. Hence, we further constructed L452R mutant viruses with or without the additional 203 mutation in the N protein and investigated these mutants in cell lines, hamsters and a human airway tissue model. We identified and validated increases in the infectivity and fitness of the L452R variants with R203M or R203K/G204R mutations relative to those without the 203 mutation. More importantly, we identified a decreased sensitivity of virus carrying the R203M N protein mutation in combination with the L452R S protein mutation towards immune serum obtained from previously SARS-CoV-2 wild type infected hamsters. Accordingly, we observed an increased fitness of the R203M/L452R virus relative to the L452R-only virus after global vaccination programmes were initiated. In summary, our results reveal that the 203 N protein mutation plays an essential role in the immune evasion and global spread of SARS-CoV-2.

**Fig. 1.**
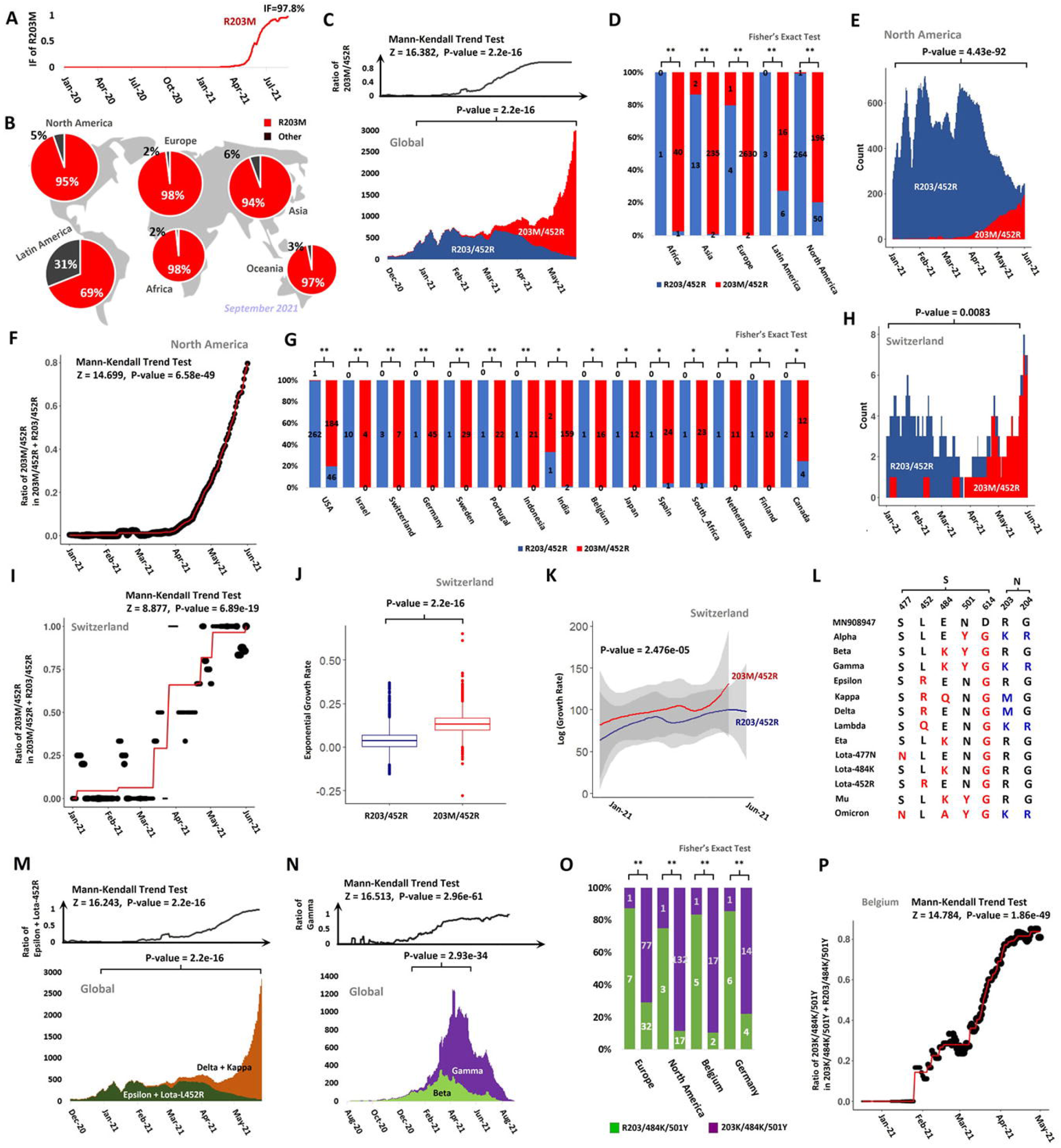
Evidence suggesting the synergistic effect between the N protein 203 mutation and L452R (A) IF changes in R203M up to September 2021. (B) The fraction of R203M on different continents in September 2021. (C) The upper panel shows the changes in the ratio of the 203M/452R in 203M/452R+R203/452R variants with the result of the Mann-Kendall trend test listed at the top. The lower panel shows the weekly running counts of the 203M/452R and R203/452R variants from December 2020 to June 2021. The result of Fisher’s exact test at the time interval bracketed is shown at the top. (D) Comparison of the fractions of the R203/452R and 203M/452R variants at two time points separated by a gap of more than 2weeks on different continents, corresponding to Table S1. ‘**’ and ‘*’ denote significance and weak significance in Fisher’s exact test, respectively. The counts of identified samples are marked within bars. (E) The comparison of weekly running counts between the R203/452R and 203M/452R variants in North America. (F) The fitted trend of the change in the fractions of the 203M/452R variant in the 203M/452R+R203/452R variants in North America with the result of Mann-Kendall trend test listed at the top. The red folded line is the maximum likelihood estimate with a nondecreasing constraint. The dot size reflects the number of identified samples on that day. (G) Comparison of the fractions of the R203/452R and 203M/452R variants in different countries, similar to (D), corresponding to Table S1. Legends in (G) follow (D). (H) The comparison of weekly running counts between the R203/452R and 203M/452R variants in Switzerland. (I) Fitted trend of the change in the fractions of the 203M/452R variant in the 203M/452R+R203/452R variants in Switzerland. Legends follow (F). (J) Comparison of the growth rates simulated in the exponential growth model between the 203M/452R (red) and R203/452R viruses (blue). (K) The growth rates (logged in the plot) over time simulated in the skygrowth coalescent model. (L) Key amino acid differences in different VOCs. The content and legends in (M) and (N) follow (C). (M) Comparison between Delta+Kappa and Epsilon+Lota-452R. (N) Comparison between Gamma and Beta. (O) Comparison of the fractions of R203/484K/501Y and 203K/484K/501Y by Fisher’s exact test at two time points, corresponding to Table S3. (P) The fitted trend of the fraction change of 203K/484K/501Y in 203K/484K/501Y+R203/484K/501Y in Switzerland like (F). Legends follow (F).

## RESULTS

### Selection advantages of the L452R variant with the nucleocapsid 203 mutation over the L452R-only variant

To evaluate the effect of R203M on Delta, we performed a comparison between the strains with both L452R and R203M and those with L452R but without R203M. We chose L452R because this S protein mutation is associated with the evasion from cellular immunity and an increase in infectivity of Delta variant ^21,26^. The comparison was performed in multiple geographically restricted contexts and in a six-month interval from the initial rise in R203M/L452R variant in January 2020 (Fig. S1C). The significance was evaluated by Fisher’s exact test of the fractions of pairs of lineages, Mann-Kendall trend test and isotonic regression analysis (for details, see the Methods section). Global, continental- and country-level analyses consistently indicate that the R203M/L452R variant has a higher growth rate than the L452R-only variant (Fig. 1C-I, Table S1, 2, IF_203M in Data S1), suggesting an increased transmission fitness of the R203M/L452R variant. To evaluate this hypothesis on a small scale, we performed a coalescent simulation for strains collected in Switzerland. Simulations with an exponential model and with a skygrowth coalescent model both confirmed the growth advantage of R203M/L452R over the L452R-only variant (Figs. 1J, K, S1D), suggesting a positive selection for the R203M/L452R mutations.

For the identified lineages, Epsilon and Lota-452R bear the L452R mutation only, while Delta and Kappa bear both R203M and L452R mutations (Fig 1L). The counts of the R203M/L452R variant are highly correlated with those of Delta + Kappa, as is the correlation between the L452R-only variant and Epsilon + Lota-452R (Figs. S1E, F, S2). Lota-452R is a sublineage of the VOC Lota bearing L452R ^27^. The overlap of Delta + Kappa and L452R/R203M (914639 strains) accounts for 82.1% of the sum-up (1113998 strains), while those of Epsilon + Lota-452R and L452R-only variant accounts for 62.2% of the sum-up. Surveys on a global scale show that the Delta and Kappa variants both have a higher growth rate than the Epsilon variant (Fig. S1G, H). Delta + Kappa variants have a higher growth rate than Epsilon + Lota-452R variants (Fig. 1M). These results confirm the selection advantage of the R203M mutation, which contributes to the superior spread of viruses carrying this mutation.

To understand whether the R203K/G204R mutations have a similar effect, we performed a comparison of the growth rate between the Beta and Gamma variants. The Beta variant bears the E484K, N501Y and D614G mutations but does not carry the R203K mutation, while the Gamma variant bears not only these three S protein mutations but also the R203K/G204R mutations (Fig. 1L). We used R203K to represent the R203K/G204R mutations here and in the following description for convenience, considering that R203K and G204R are completely linked ^12^. Our results show that the Gamma variant have a higher growth rate than Beta (Fig. 1N). Thereafter, we compared the growth rate between the 203K/484K/501Y variant and the R203/484K/501Y variant at different geographical levels. We did not consider D614G in this comparison because this mutation has been fixed in virus genomes around the world (Fig 1L) since July 2020 ^23,28^. In line with the results of the comparison results between the Gamma and Beta variants, the growth rate of the 203K/484K/501Y variant was consistently higher than that of the R203/484K/501Y variant (Figs. 1O, P, S1I, Table S3, 4 and IF_203K in Data S1). These results suggest a synergistic effect between the N protein 203 mutation and the functional S protein mutations.

### A synergistic effect of the N protein 203 mutation on the S protein L452R mutation

To evaluate the synergistic effect between the N protein 203 mutation and the S protein 452 mutation, we constructed four SARS-CoV-2 variants, namely, L452R (without the N protein 203 mutation), R203M (without the S protein L452R mutation), R203M/L452R, and R203K/L452R, based on the USA_WA1/2020 SARS-CoV-2 sequence (GenBank accession No. MT020880) by using a reverse genetic method and testing viral replication in hamsters. Three- to four-week-old hamsters were anaesthetized with isoflurane and infected intranasally with a 1:1 mixture of wild type or mutant virus. The quantity of released virus was assessed in a competition experiment at 4 days and 7 days after infection (Fig. 2 and S3). As a result, a higher ratio of the R203M/L452R variant to the L452R variant, as well as the R203K/L452R variant to the L452R variant was observed at 4 days after infection (Fig. 2B, C), indicating a replication advantage of the viruses bearing the nucleocapsid (N) and spike (S) joint mutations in hamsters. We then compared the competition between the R203M and R203K variants with or without the presence of the L452R mutation. R203M-only variant exhibited increased viral replication (Fig. 2A). However, regardless of whether L452 was mutated, there was no significant difference in viral competition between the R203M and R203K variants (Fig. 2D, E). The ratios of the R203M/L452R and R203K/L452R variants to the L452R variant were both greater than 1 at 4 and 7 days postinfection (dpi) (Fig. S3B, C), indicating that the joint mutations of R203M/L452R or R203K/L452R have a consistent replication advantage over the L452R mutation alone in hamsters.

**Fig. 2.**
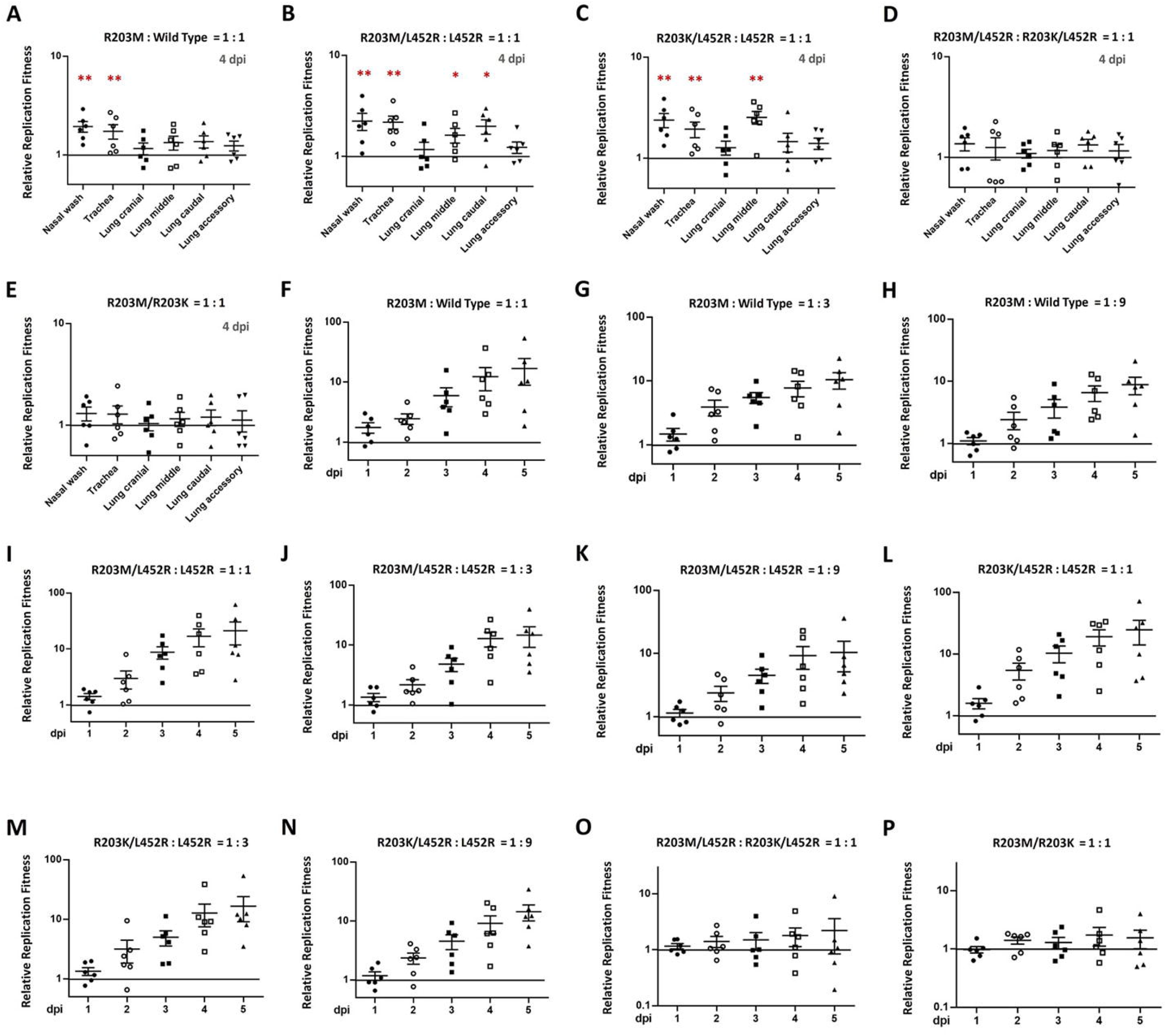
Experimental results to validate the synergistic effect between N and S mutations. (A-E) Hamsters were intranasally infected with a 1:1 mixture of wild-type or mutant viruses as indicated. Nasal washes, tracheas and lungs were collected on day 4 after infection. The relative RNA amounts of R203M: wild type (A), L452R/R203M: L452R (B), L452R/R203K: L452R (C), L452R/R203M: L452R/R203K (D), and R203M: R203K (E) were assessed by RT-PCR and Sanger sequencing. Log10 scale was used for the Y-axis. Data are presented as the mean ± s.e.m.. Dots represent individual hamsters (n = 6). *, p<0.05, **, p<0.01. (F-P) Mixtures of wild type or mutant viruses with an initial ratio of 1:1 (F, I, L, O, P), 1:3 (G, J, M) or 1:9 (H, K, N) were inoculated into human airway tissue cultures at a MOI of 5 as indicated. Viral ratios of R203M: wild type (F-H), R203M/L452R : L452R (I-K), R203K/L452R : L452R (L-N), R203M/L452R : R203K/L452R (O) and R203M: R203K (P) from 1-5 dpi were measured by RT-PCR and Sanger sequencing. Log10 scale was used for the Y-axis. All data in (F-P) are represented as the mean ± s.e.m.. Dots represent individual hamsters (n = 6).

Furthermore, we performed competition experiments using a human airway tissue culture model (Fig. 2F-P). After infecting the airway culture with wild-type and mutant viruses at a ratio of 1:1, the ratios of the L452R viruses bearing the N protein mutation to the L452R-only virus increased from 1 to 5 dpi (Fig. 2I, L). After infecting the tissues with a 1:3 or 1:9 ratio of the R203M/L452R virus to the L452R-only virus and the R203K/L452R virus to the L452R-only virus, the joint mutants rapidly overcame their initial deficit and showed an advantage over the L452R variants (Fig. 2J, K, M, N). All these results indicate that the joint mutations R203M/L452R or R203K/L452R in SARS-CoV-2 rapidly outcompeted the L452R-only variant in both hamsters and human airway tissues, suggesting a synergistic effect of the N protein mutation and the S protein mutation. We observed an alteration in the polymerization state of the N protein after the 203 mutation (Fig. S4, for details, see the Methods section), which may cause the synergistic effect.

### R203+L452R joint mutants show higher infectivity than the L452R-only variant

We compared the infectivity of the constructed variants in the Vero E6 monkey kidney cell line and Calu-3 human lung epithelial cell line. The results show that all the variants replicated with higher extracellular viral RNA production than the wild type at 36 h post-infection (Fig. 3A). The infectivity of the virus was measured by PFU titres and viral subgenomic RNA (E sgRNA) loads. As a result, similar trends of infectious titres in the variants were observed. Specifically, the difference between the production of PFU titres in the R203+L452R joint mutants (R203M/L452R or R203K/L452R) and the wild type, is more significant than that between the L452R-only mutant and the wild-type virus (P-value < 0.01 versus P-value < 0.05; Fig. 3B). However, we found that there were no significant differences in the E sgRNA loads among the variants (Fig. 3C). In the Calu-3 cells, the difference between the production of PFU titres/E sgRNA loads in the R203+L452R joint mutants and the wild type was more significant than that between the L452R-only mutant and the wild type; however, this phenomenon was not found in the extracellular viral RNA levels (Fig. 3D-F). These results indicate that the N protein 203 mutation further enhances the viral replication and infectivity of L452R virus.

**Fig. 3.**
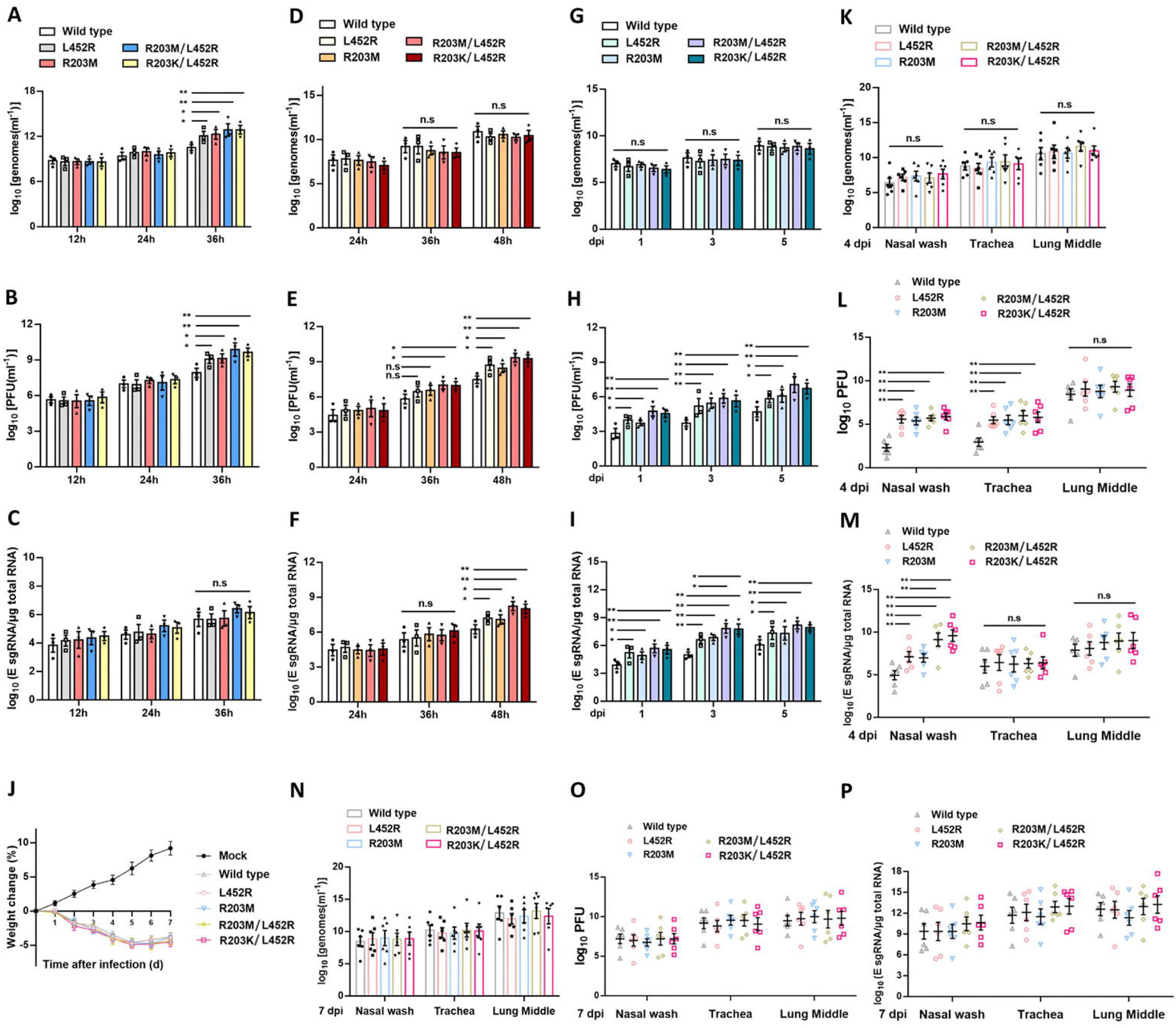
Comparison of the infectivity between the virus with R203+L452R joint mutations and that with L452R alone by cell and animal experiments. (A-F) Viral replication, infectious viral titres, and viral subgenomic RNA of wild type or mutant viruses produced from Vero E6 (A-C) and Calu-3 (D-F) cell cultures were measured. Cells were infected at a MOI of 0.01. Genomic RNA levels (A, D), PFU titres (B, E) and E sgRNA loads (C, F) were determined by plaque assays and qRT-PCR, respectively. (G-I) Wild-type or mutant viruses were inoculated into primary human airway tissues at an MOI of 5. After incubation for 2 h, the cultures were washed with PBS and maintained for 5 days. Genomic RNA levels (G), PFU titres (H) and E sgRNA loads (I) were determined by plaque assays and qRT-PCR, respectively. Data are represented as the mean ± s.e.m.. *, p<0.05, **, p<0.01. Abbreviation: n.s., nonsignificant. (J-M) Hamsters were intranasally infected with 2×10^4^ PFU of the wild type or mutant viruses as indicated. Weight loss (J) was monitored for 7 days. Data are presented as the mean ± s.e.m.; n = 12 (all cohorts) at days 0–4; n = 6 (all cohorts) at days 5–7. Weight loss was analysed by two-factor analysis of variance (ANOVA) with Tukey’s post hoc test. Viral genomic RNA levels (K), PFU titres (L) and E sgRNA loads (M) were quantified in nasal wash, trachea and lung samples at 4 dpi. Viral genomic RNA levels (N), PFU titres (O) and E sgRNA loads (P) were also quantified in nasal wash, trachea and lung samples at 7 dpi. Dots represent individual hamsters (n = 6). Data are presented as the mean ± s.e.m.. **, p<0.01. Abbreviation: n.s., nonsignificant.

In the comparison conducted in a human airway model, we found no differences in viral RNA yields between the variants (Fig. 3G). However, the PFU titres and viral E sgRNA loads of all mutants are significantly higher than those of the wild-type virus (Fig. 3H, I). In particular, it was observed that the E sgRNA loads in the R203+L452R joint mutants were significantly higher than those in the L452R virus at 3 dpi (Fig. 3I). A similar comparison was performed in hamsters. Hamsters were anaesthetized with isoflurane and infected intranasally with a total of 2×10^4^ PFU of the variants or wild-type virus. Hamsters infected with different viruses exhibited similar weight loss (Fig. 3J). At 4 dpi and 7 dpi, the four variants and the wild-type virus produced a nearly identical level of viral RNA across all organs (Fig. 3K, N). We further compared the infectivity of the variants by measuring PFU titres and E sgRNA loads. The results show that the infectious viral titres measured in nasal wash and trachea samples and the E sgRNA loads measured in nasal wash samples from hamsters infected with the four variants were consistently higher than those from hamsters infected with the wild type virus at 4 dpi (Fig. 3L, M), but not at 7 dpi (Fig. 3O, P). Similar to our previous observations in a human airway model, the E sgRNA loads of the R203+L452R joint mutants were significantly higher than those of the L452R virus in nasal wash samples (Fig. 3M). Taken together, these results demonstrate that the N protein 203 mutation further increased the viral replication efficiency and virion infectivity of the L452R variant in both a human airway model and hamsters.

### The 203 mutation enhances the neutralization resistance of the L452R virus

We next evaluated the effect of the 203 mutation on the sensitivity of the mutant viruses to neutralization serum. Neutralization titres for a panel of sera collected from hamsters infected with the wild-type virus were analysed using the L452R, R203M, L452R/R203M and L452R/R203K variants containing an mNeonGreen reporter, respectively. In comparison with those for the wild-type virus, the neutralization titres for the L452R virus in all the sera were reduced by 1.24- to 3.05-fold (mean 1.90-fold) (Fig. 4A, B). In contrast, the neutralization titres for the R203M mutation group increased by 1.16- to 2.22-fold, with a mean of 1.49-fold (1/NT_50_: 0.45- to 0.86-fold, mean 0.67-fold) (Fig. 4A, B), which is consistent with our previous finding that the N protein 203 mutation confers increased susceptibility of SARS-CoV-2 to serum neutralization ^23^. Interestingly, further reduced neutralization titres for the joint mutants L452R/R203M and L452R/R203K were found in the sera collected from hamsters, with 1.65- to 5.52-fold (mean 3.28-fold) and 1.71- to 5.07-fold (mean 3.07-fold) reductions when compared with those for the wild-type virus (Fig. 4A, B). The lowest neutralization titres for the L452R, L452R/R203M and L452R/R203K variants were found in serum 5 (Fig. 4C), while the highest neutralization titres for the R203M variant were found in serum 6 (Fig. 4D). These data suggests that the L452R mutation confers the escape of SARS-CoV-2 variants from antibody neutralization and that the N protein 203 mutation further enhances the escape phenotype.

**Fig. 4.**
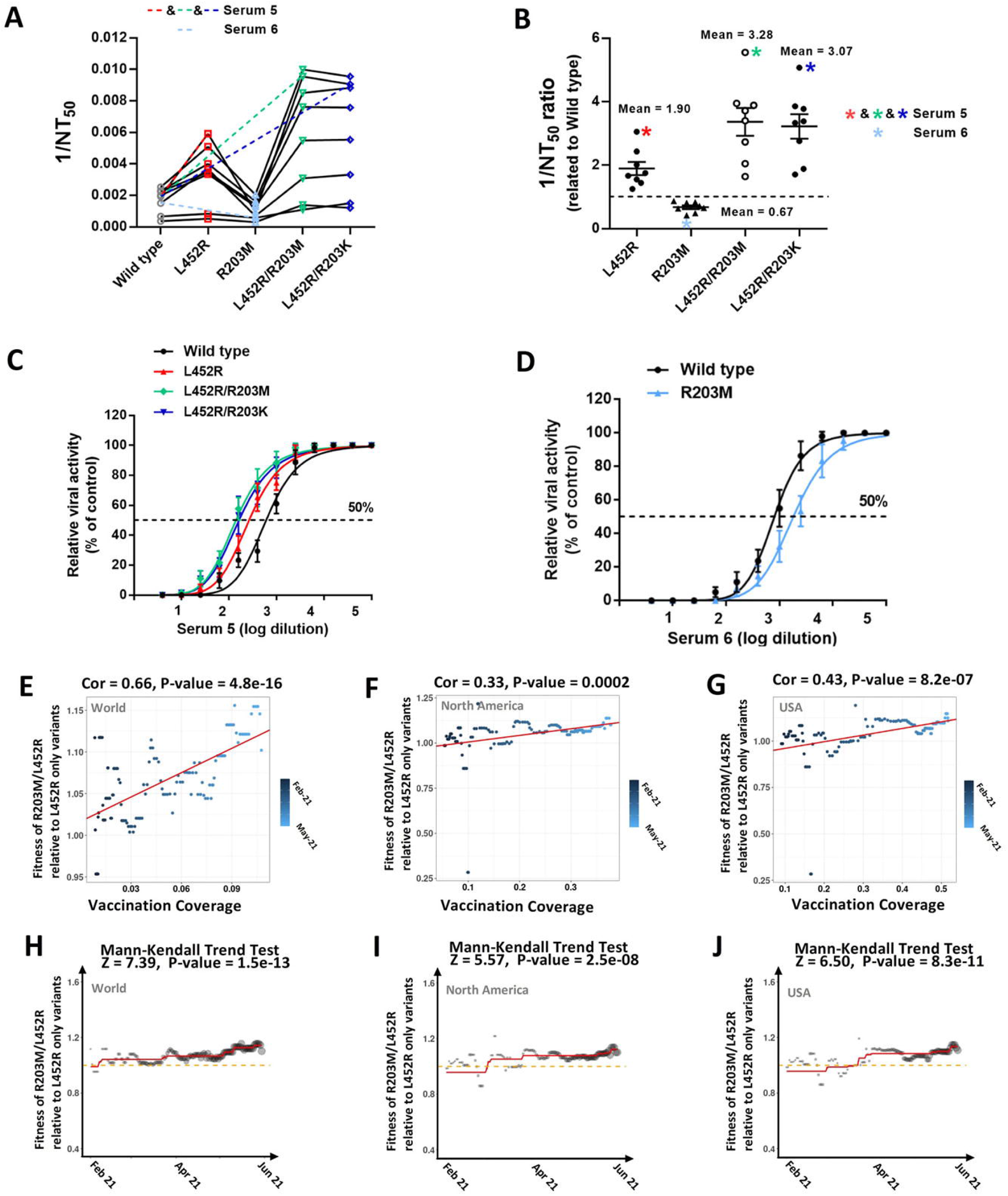
Evidence that demonstrates the contribution of the 203 mutation to immune evasion. (A) Neutralization assay of hamster sera against the L452R, R203M, L452R/R203M and L452R/R203K variants containing an mNeonGreen reporter. 1/NT_50_ values were plotted. Symbols represent sera from individual hamsters (n = 8). (B) 1/NT_50_ values of L452R, R203M, L452R/R203M and L452R/R203K variants to wild type virus were calculated. Data are represented as the mean ± s.e.m.. Symbols represent sera from individual hamsters (n = 8). (C-D) Neutralization curves of serum 5 (C) and serum 6 (D) from individual hamsters. The solid line represents the fitted curve, and the dotted line indicates 50% viral inhibition. (E-G) Correlation analysis results between the fitness of R203M/L452R relative to the L452R-only variant (y-axis) and VC (x-axis) in the world (E), in North America (F) and in the USA (G). The colour of the dots corresponds to the date. (H-J) The fitted trend of the fitness of the R203M/L452R variant relative to the L452R-only variant in the world (H), North America (I) and the USA (J). The results of the Mann-Kendall trend test are listed at the top.

### Increased fitness of the R203M/L452R variant relative to L452R-only variant after the initiation of global vaccination programmes

We evaluated the changes in the fitness of the R203M/L452R variant relative to that of the L452R-only variant and the correlation between the fitness and vaccination coverage (VC). The relative fitness was calculated based on the changes in the IF of the R203M/L452R and L452R-only variants. The evaluation was performed at hierarchical geographical levels. We observed a continuously increasing relative fitness of the R203M/L452R variant, and the increase is significantly correlated with the increase in VC (Fig. 4E-J, Table S5 and RF_203M452R_to_203R452R in Data S1). The increase in relative fitness and the positive correlation is likely due to the elevated resistance to neutralization of the R203M/L452R variant. In the same way, we also evaluated the fitness of the R203M/L452R variant (Delta) relative to the R203K/N501Y variant (Alpha). There is an upward trend of the relative fitness of the 203M/452R variant and the trend is significantly correlated with the values of VC (Fig. S5A-F, Table S6 and RF_203M452R_to_203K501Y in Data S1). These results may explain the fast replacement of Alpha by Delta and the low IF of Delta for a lone time (Figs. S1A, S5H). The 203M/452R and Delta variants appeared as early as March 2020 and May 2020 ^29^, respectively (Fig. S5G, H). Global vaccination programmes may have boosted the expansion of the R203M/L452R and Delta variants (Fig. S5I), considering that vaccinated people may be more vulnerable to Delta infection than the infection of other VOCs ^30^.

### The synergistic effect of R203M and L452R on disease severity is not significant

We compared the inflammatory damage in the lungs of hamsters infected with wild-type and mutant viruses. The results show that hamsters infected with the four variants (R203M, R203K, R203M/L452R and R203K/L452R) displayed more extensive inflammatory damage and pulmonary vascular congestion than hamsters infected with the wild-type virus (Fig. 5A-F). However, we did not observe a significant difference in pathology scoring between the joint mutants and the L452R-only variants (Fig. 5F). In a global survey based on the documented clinical information of strains from the Global Initiative on Sharing Avian Flu Data (GISAID), we also did not find a consistently increased virulence of the R203+L452R variant relative to the L452R-only variant (Fig. 5G).

**Fig. 5.**
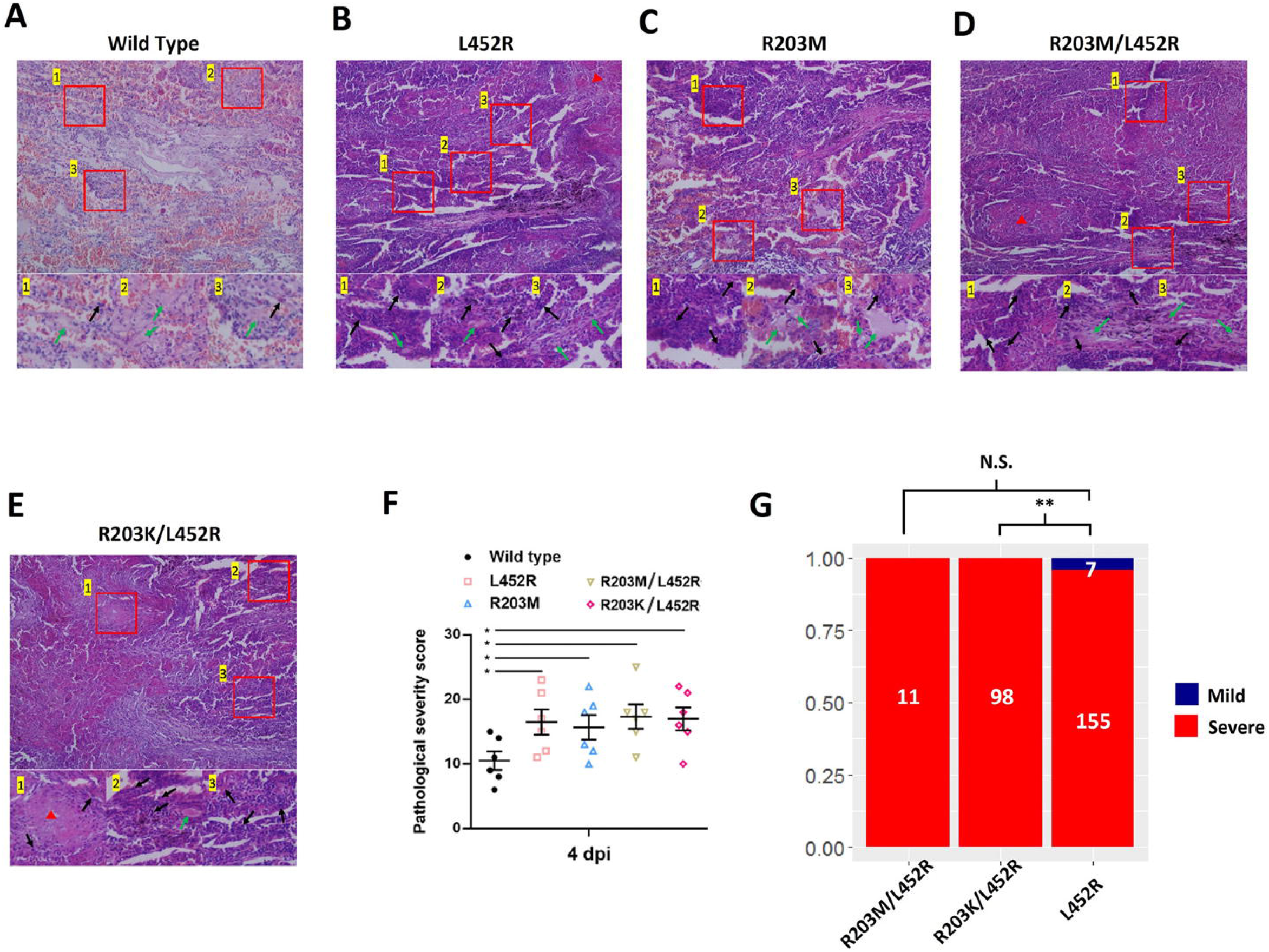
Evaluation of the effects of the 203 mutation on L452R virulence. (A-F) Haematoxylin and eosin (H&E) staining of lung sections collected at 4 dpi from hamsters infected with wild-type (A), L452R (B), R203M (C), R203M/L452R (D) or R203K/L452R (E) variants. The lower photographs are higher magnification images of the regions denoted by rectangles in the upper photographs. The upper panel shows inflammatory damage with vascular congestion. The lower panel shows bronchioles with aggregation of inflammatory cells (black arrow) and surrounding alveolar wall infiltration (green arrow). Red arrowheads indicate the alveolar parenchymal lesions. Scale bar = 100 μm. (F) Histopathology score of lung sections. Lung lobes were scored individually using the scoring system described in the methods. The scores of 5 slices from each hamster were added to determine the total pathology score per animal. Dots represent individual hamsters (n = 6). Data are presented as the mean ± s.e.m.. *, p<0.05. (G) Prediction of the clinical outcomes of R203M/L452R, R203K/L452R and L452R. The Y axis shows the ratios between lineages with a clinical status, mild or severe. Lineage numbers are provided within bars. The significance was tested by Chi-squared test. N.S. denotes not significant.

## DISCUSSION

Genetic mutation has produced multiple SARS-CoV-2 variants with increased transmissibility, infectivity and immune escape. Commencing from March 2021, the SARS-CoV-2 Delta variants rapidly replaced the previously dominant variants, including Alpha, which is ∼50% more transmissible than the Wuhan-Hu-1 strain ^16^. The emergence of Delta after the initiation of global vaccination programmes indicates the importance of identifying the dominant mutation that contributes to immune evasion and enhanced infectivity. This is essential for the development of vaccines and medicines. Our previous work reported the rapid spread of the R203K/G204R mutation ^12^ and further adaptiveness of the mutation ^23^. In our consistent tracking of the mutations occurring in VOCs, we noticed that the newly evolved dominant lineage Delta possesses an R203M mutation in the N protein instead of the R203K/G204R mutations. Through statistical analyses of Delta and comparisons between Delta and other VOCs, we observed an association between the occurrence of Delta and immune escape in vaccinated patients. Furthermore, we found a synergistic effect of the S protein mutations and the N protein mutations and a correlation between the combinational mutations R203M/L452R and the resistance of Delta variant to post vaccination immunity.

Based on these findings, subsequent virological experiments using the mutant virus were performed. The results of our experiments show that the 203M virus exhibits increased replication in cell lines and animal models, similar to the phenotype of the 203K/204R virus. Consistent with our previous findings in the 203K/204R virus ^23^, the 203M-only virus has an increased sensitivity to neutralizing serum (Fig. 4A, B), suggesting a similar functional output of the 203 mutations, even though we could not explain the mechanism of the decreased sensitivity to the neutralizing sera introduced by the N protein mutation. Surprisingly, the outcome was reversed for the 203 mutations in combination with the S protein mutation L452R. The 452R/203M double mutant virus shows dramatically decreased sensitivity to neutralizing sera compared with the 452R single-mutant virus. The experimental synergistic effect of the S and N mutations is well consistent with our statistical observation of the emergence of Delta variants. The reason for the sensitivity to the neutralizing sera of the mutant virus is a mystery and needs more in-depth work in the future.

There are several hypotheses to explain the increased transmissibility of Delta. Most studies have focused on S protein mutations, such as P681R and L452R. P681R leads to more efficient furin cleavage ^31^, and L452R decreases the sensitivity to antibody neutralization ^21^. The escape from antibody neutralization of Delta is regarded as associated with the S protein mutation only. Although there are several mutations in the S protein, there are mutations in the N protein (e.g., R203M) fixed early in the emergence of different VOCs, e.g., Delta (Fig. S5K). Our results highlight the essential roles of N protein mutations, in addition to S protein mutations, in the immune evasion of SARS-CoV-2. The rapidly fixed mutation R203M in Delta variants, which is in the same location as the R203K mutation that occurs in Alpha variant, is likely not a coincidence. The improved mRNA packaging and luciferase induction associated with the R203M mutation based on the virus-like particle (VLP) assay also confirmed these results, although the mechanism of the N protein mutations is attributed to a previously unknown strategy ^25^. The 203 mutation may influence the polymerization, phase separation or phosphorylation of the N protein and further impact the assembly of ribonucleocapsid (RNP). The influence not only increases the fitness of the virus but also unexpectedly enhances immune evasion. The reduced vaccine effectiveness resulting from the R203M mutation further causes the escape of the Delta variants from the neutralizing antibody elicited by the previous vaccination against SARS-CoV-2. The N protein mutation should be considered a main factor contributing to Delta dominance.

Our results demonstrate that the dominantly emerging Delta variants, which escape the immunity associated with current global vaccination programmes, are essentially facilitated by the R203M mutation in the N protein. These findings highlight the important role of N protein mutations in persistently circulating SARS-CoV-2 variants. In addition, our results prove the synergistic effect of S and N protein mutations on the increased immune evasion of SARS-CoV-2. A newly reported VOC Omicron^32,33^ also contains the 203 mutation, R203K/G204R (Fig. 1L), suggesting a potential universal effect of the specific mutation. Persistent and close attention to the 203 mutation in Omicron and subsequent emerging variants is essential moving forward.

### Limitations of the study

The limitations of our work are mostly attributed to the lack of mechanistic studies of the N protein mutation. We demonstrated a positive contribution of the R203M mutation to the transmission and resistance to neutralization of Delta. However, the clarification of the detailed mechanism requires more extensive biochemical and structural research. Detailed biochemistry and structural studies would be helpful to characterize the biological function of the N mutation. We did not observe an increased virulence associated with the R203M/L452R variant relative to that of the L452R-only variant. These results do not indicate that there is no contribution of the R203M mutation to the disease severity of Delta. Hence, a more comprehensive survey and in-depth investigation are needed.

## ACKNOWLEDGMENTS

We gratefully acknowledge the submitting and originating laboratories where genetic sequence data were generated and shared via NCBI and the GISAID Initiative. This work was supported by grants from the National Natural Science Foundation of China, SGC’s Rapid Response Funding for COVID-19 (C-0002), the Fundamental Research Funds for the Central Universities (2021CDJZYJH-002, 2021CDJYGRH-009), the National Key Research and Development Program (2019YFC1604600), the National Natural Science Foundation of China (81970008, 32170661, 32100156 and 82000020), the Youth Innovative Talents Training Project of Chongqing (CY210102), the Chongqing Talents: Exceptional Young Talents Project (cstc2021ycjh-bgzxm0099) and the National Natural Science Foundation of Hebei Province (19226631D).

## AUTHOR CONTRIBUTIONS

Z.Z., G.M, Y.X., Y.Z., K.M. and W.T. collected the data, performed population genetic analyses and took part in the editing of the manuscript. X.L., W.X., N.X., B.F., J.T., D.K. and H.W. performed the virology experiments. G.M., Y.Z. and S.S. performed the protein structure and sequence analysis. Z.Z., G.M. and H.W. conceived the idea. Z.Z., G.M., H.W., J.T., D.K. and X.L. wrote the manuscript. Z.Z., G.M. and H.W. coordinate the project.

## Materials and Methods

### Evolutionary analyses of the strains with and without the N protein 203 mutation

The source data includes 3438667 full-length genomic sequences of SARS-CoV-2 collected from December 2019 to September 2021 (SARS2_List.xls in Data S1), downloaded from GISAID (www.gisaid.org), NCBI and CoVdb ^34^. We performed pairwise alignments of the 3438667 SARS-CoV-2 genomic sequences and the genomic sequence of MN908947 collected on December 2019 ^1^ and considered these sequence as the reference sequence in this study. Based on the alignments, we identified the mutations in the genome of each strain and performed statistical analysis of the R203M mutation. We wrote Perl scripts to count the weekly running counts of the strains with the R203M/L452R mutation and those with the L452R mutation only. To evaluate the changes in IF of the R203M/L452R variant relative to those of the L452R-only variant, we performed Fisher’s exact test of the fraction of pairs of lineages at the onset, when there was an introduction of a new variant, and on the day after more than two weeks. For the limited number of L452R only variants, we used loose criteria. We required that the counts were higher than zero for both lineages at the two time points. Furthermore, we built a maximum likelihood estimation line of the fraction trend and evaluated the trend by the Mann-Kendall trend test and isotonic regression analysis. The evaluation was performed at three hierarchical geographic levels (world, continent and country). We did not perform a test at the city level for the limited number of the L452R-only variant. We also performed binomial tests at the continent level and the country level. We used the same method to evaluate the growth rates between the R203/484K/501Y and 203K/484K/501Y variants. According to the annotation from GISAID, we identified the mutations of VOCs, including Alpha, Beta, Gamma, Epsilon, Eta, Lambda, Mu, Kappa, Delta and Omicron. For example, we identified 31 mutations in Delta. For Lota, we identified the mutations based on published work ^27^, according to which we attributed the Lota variants with the L452R mutation to Lota-452R, those with S477N to Lota-S477N and those with E484K to Lota-484K. Based on the mutation information of the lineages, we performed statistics of the change in IF for VOCs. Based on previous work^23,35^, we evaluated the transmission advantages of the R203M/L452R variant relative to those of the L452R-only variant in Switzerland. We performed simulations of the strains from January 2021 to June 2021 in an exponential growth model by using BEAST v1.10.4 ^36^ with an HYK substitution model and a strict clock type and a chain length of 1000000. We used FastTree ^37^ (the parameters are -gtr -nt -boot 1000) to build the phylogenetic tree of the R203M/L452R and L452R variants, respectively. Then, we performed phylodynamic inference of effective population size for time-scaled phylogenies by the R package Skygrowth ^38^ with parameters res = 300 and tau0 = 0.1. We performed analyses of the simulation data and built figures with the R packages gdata, ggplot2 and dplyr.

### Preparatory work of animals, cells and infection

Six- to eight-week old female golden Syrian hamsters were obtained from Vital River Laboratories (Beijing, China). All animal experimental procedures were approved by the Animal Ethics Commission of the School of Life Sciences, Chongqing University. The hamsters were anaesthetized by isoflurane and infected intranasally with 2×10^4^ PFU virus. Specifically, 12 hamsters received wild-type virus, 12 received L452R mutant virus, 12 received R203M mutant virus, 12 received L452R/R203M mutant virus, 12 received L452R/R203K mutant virus, and 12 received phosphate-buffered saline (PBS) (Mock). For the competition experiment, 12 hamsters received a 1:1 mixture of the R203M mutant and wild-type virus, 12 received a 1:1 mixture of the L452R/R203M and L452R viruses, 12 received a 1:1 mixture of the L452R/R203K and L452R viruses, 12 received a 1:1 mixture of the L452R/R203M and L452R/R203K viruses, and 12 received a 1:1 mixture of the R203M and R203K viruses. The infected hamsters were weighed and recorded daily. On the 4th and 7th days after infection, cohorts of 6 infected hamsters were anaesthetized with isoflurane, and nasal washes were collected with sterile dulbecco phosphate-buffered saline (DPBS). Immediately after nasal wash, the hamsters were humanely euthanized, and the trachea and lung middle lobe were obtained as previously described ^11^. All hamster operations were performed under isoflurane anaesthesia to minimize animal pain. Human lung adenocarcinoma epithelial Calu-3 cells (ATCC) and African green monkey kidney epithelial Vero E6 cells (ATCC) were maintained at 37⍰°C with 5% CO_2_ in high-glucose Dulbecco’s modified Eagle’s medium (DMEM, Gibco) supplemented with 10% FBS (Gibco). Cells were infected with a multiplicity of infection (MOI) of 0.01 at the indicated time points. All live virus experiments were carried out under biosafety level 3 (BSL3+) conditions.

### Generation of the SARS-CoV-2 mutant viruses

A reverse genetic method was used to generate the wild-type SARS-CoV-2 (USA_WA1/2020 SARS-CoV-2 sequence, GenBank accession No. MT020880) virus, as previously described ^11,39,40^. Using standard molecular cloning methods, seven different DNA fragments spanning the entire genome of SARS-CoV-2, were synthesized by Beijing Genomics Institute (BGI, Shanghai, China) and cloned into the pCC1 or pUC57 plasmid. The sequences of the F1∼F7 fragments as well as the restriction enzymes for digestion and ligation were described in our previous work ^23^. Full-length cDNA was assembled, and recombinant SARS-CoV-2 virus was recovered as previously described ^11,39,40^. Specifically, the full-length cDNA of SARS-CoV-2 was assembled through the *in vitro* ligation of contiguous cDNA fragments. Then, full-length genomic RNA was obtained via *in vitro* transcription, followed by electroporation into Vero E6 cells. The SARS-CoV-2 virus was harvested at 40 h post-electroporation. A plaque assay was performed to determine viral titres. The generation of the L452R, R203M, L452R/R203M, and L452R/R203K variants was performed by overlap-extension PCR. The mutation sites and primers are shown in Table S7.

### Plaque assays and neutralization assays

Plaque assays were performed following our previous work ^23^. Briefly, approximately 1×10^6^ cells were seeded into 6-well plates, followed by culture in 5% CO_2_ at 37⍰°C for 12 h. Wild type or mutant viruses were serially diluted in DMEM containing 2% FBS, and 200 μL aliquots were added to the cells. The cells were co-incubated with viruses for 1 h and supplemented with overlay medium containing DMEM with 2% FBS and 1% Sea-Plaque agarose. After 2 days of incubation, neutral red was used to stain the plates, and plaques were measured in a light box.

Neutralization assays were performed using the L452R, R203M, L452R/R203M, and L452R/R203K viruses containing an mNeonGreen reporter, as previously described ^11,23^. Briefly, Vero E6 cells were plated in 96-well plates. The next day sera were serially diluted and mixed with mNeonGreen viruses at 37⍰°C for 1 h incubation. The virus-serum mixture was then transferred to a cell plate at a final MOI of 2.0. After 20 h, cell nuclei were stained with Hoechst 33342 solution, and mNeonGreen fluorescence was quantified. The numbers of mNeonGreen-positive cells and total cells were measured in each well. The infection rate was determined by dividing the number of mNeonGreen-positive cells by the total number of cells. The relative infection rate was obtained by normalizing the infection rate of the serum-treated groups to that of the non-serum-treated control groups. The fold dilution that neutralized 50% of mNeonGreen fluorescence (NT_50_) was determined using a nonlinear regression method. GraphPad Prism 8 was used to plot the curves of the relative infection rates versus the serum dilutions (log_10_ values).

### Viral infection in a primary human airway tissue model

The wild-type virus or the variants were inoculated into a primary human airway tissue culture at an MOI of 5. After 2 h of infection, the inoculum was removed, and the culture was washed with PBS three times. The infected epithelial cells were maintained in the apical well without any medium, and the medium was provided to the culture through the basal well. From day 1 to day 5, 300 μL PBS was added to the apical side of the airway culture, followed by incubation at 37⍰°C for 30 min to elute the released viruses.

### Competition, viral subgenomic RNA and genomic RNA assays

Competition assays were performed by RT-PCR with quantification of Sanger peak heights, as previously described ^23^. In detail, the wild-type virus and the variants were mixed at the indicated ratios based on their PFU titres. A pair of common primers (SARS-CoV-2 28354F and SARS-CoV-2 28949R, Table S8) was used to quantify the R203M: wild type, L452R/R203M: L452R, L452R/R203K: L452R, L452R/R203M: L452R/R203K and R203M: R203K ratios. Using a SuperScript III One-Step RT–PCR kit (Thermo Fisher Scientific), the RT-PCR product (596 bp) was amplified from the extracted RNA according to the manufacturer’s instructions. A GeneJET PCR Purification kit (Thermo Fisher Scientific) was used to purify the PCR product, which was submitted to Sanger sequencing (BGI, Shanghai, China) (Primer for Sanger sequencing, Table S8). The sequence electropherograms were then scored with a QSV analyser to determine the proportion of different viruses.

A viral subgenomic RNA assay was performed with a leader-specific forward primer, a reverse primer and a probe that targets the envelope (E) protein gene sequences, as previously described ^23,41^. Infectious cell lysates were harvested at the indicated time points, followed by total RNA extraction with an RNeasy Mini kit (QIAGEN, Hilden, Germany). RT-PCR was performed using a SuperScript III One-Step RT–PCR kit (Thermo Fisher Scientific) and an ABI StepOnePlus PCR system (Applied Biosystems, CA, USA), according to the manufacturer’s instructions. The primers were E_Sarbeco_F, E_Sarbeco_R and E_Sarbeco_P1 (Table S8), as previously described ^23,41^. The Orf1ab gene was used for the quantification of viral genomic RNA. The primers were SARS-CoV2.ORF1ab.F, SARS-CoV2.ORF1ab.R and SARS-CoV2.ORF1ab.P, as previously described ^23,42^.

### Pathological examination

The hamsters were anaesthetized with isoflurane and tissues were harvested at the indicated time points. Tissues were fixed in 10% formalin, trimmed and embedded in paraffin. The paraffin blocks were sectioned into a thickness of 3 μm for haematoxylin and eosin staining. The histopathology score of the infected hamsters was based on the percentage of the area of inflammation in each section of the lung lobe obtained from each animal, using a semi-quantitative pathology scoring system, as previously described ^23^. 0, no pathological change; 1, affected area (≤10%); 2, affected area (>10% and ≤30%); 3, affected area (>30% and ≤50%); and 4, affected area (>50%). Five sections were collected from each hamster, and the scores of lung sections were added to determine the total pathology score of each animal. All scoring was performed by the same operator to ensure consistency of scoring.

### The inference of relative fitness based on changes in IF of the variants

According to the theory of classical population genetics ^43^, for two genotypes A and B in a population, the change in proportion (Δp) is:

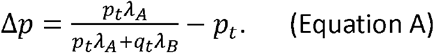

Here, *p*_*t*_ and *q*_*t*_ are the proportions of A and B in the population at moment t, respectively. *λ*_*A*_ and *λ*_*b*_ are the absolute fitness of A and B, respectively. We manipulated the equation and obtained:

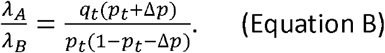

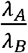 is the fitness of A relative to B at moment t. Based on Equation B, we inferred the relative fitness of the R203M/L452R variant relative to the L452R-only variant. For the sufficiency of samples, we required that the average count per day be higher than 10 for both lineages. For these, the time interval was from February 1^st^, 2021, one month post the initial rise in the R203M/L452R variant, to May 30^th^, 2021, when the IF of the L452R-only variant was decreasing but the number of L452R-only samples was still sufficient for statistics. We counted the weekly median of the relative fitness per day. We obtained the records of vaccination coverage (VC, people vaccinated per hundred) in different regions from Our World In Data (ourworldindata.org) and performed a correlation test between the weekly median of the relative fitness and VC at different geographic levels. For the limited number of identified L452R samples, we did not perform statistics on a smaller scale. We also built a maximum likelihood estimation line of the fitness trend and evaluated the trend by the Mann-Kendall trend test. In the same way, we evaluated the relative fitness of the R203M/L452R variant relative to the R203K/N510Y variant. For the sufficiency of samples, we chose the time interval from February 1^st^, 2021 to Aug 30^th^, 2021 (Fig. S5J). We kept the results with statistical significance and performed a binomial test at each geographic level.

### Function prediction based on clinical data

We manually gathered the clinical information of 115032 SARS-CoV-2 strains (Clinical_Info.xls in Data S1) from GISAID. Following the pipeline in our previous work ^12^, we grouped this information into a pair of opposite patient statuses, mild and severe, according to a series of keywords (Table S9). To avoid the impact of other mutations, we clustered strains with the same sequence in the mutation sites of Delta, excluding the S protein mutation 452 and N protein mutation 203, into groups and selected the groups with more than 20 strains. We performed multiple alignment of the consistent sequences of these groups and built a phylogenetic tree. Based on the tree we chose the groups akin to each other. We used the strains of the selected groups to perform functional prediction. We compared the fractions of mild and severe cases and tested the significance by the Chi-squared test.

### Expression and purification of the N protein

The constructed expression cassette carrying protein-encoding genes (N-native, N-R203M and N-R204K) (with Strep-tag) was integrated into yeast competent cells to give a new expression strain as previously described ^44^. Transformants were cultured on SCD agar plates plate at 30 °C for 2 days. Then newly expressed strains were transferred to YPD for propagation and protein expression. The culture was harvested by centrifugation at 6000 rpm for 10 min when the OD600 reached approximately 3.0. Cell pellets were resuspended in 50 mL of PBS (pH 7.5), followed by cell disruption by utilizing a homogenizing machine. The supernatant of the cell disruption product was collected after centrifugation at 17000 rpm for 90 min, and all the supernatant was loaded onto a Strep-Tactin XT gravity-flow column (IBA). The collected eluents were analysed with SDS-PAGE, and the final elution was selected based on purity. The final pool was concentrated to > 1 mg/mL using a Merck-Millipore 5 kDa, 15 mL concentrator. The BCA assay was performed on microtiter plates to determine the N protein concentration.

### Size exclusion chromatography (SEC) analysis

SEC analysis of purified N protein was performed by Superdex 200 16/600 (GE healthcare) at 4 °C on an AKTA system. The mobile phase consisted of PBS with a flow rate of 1 mL/min. The eluent of each peak was gathered and analysed by SDS-PAGE.

### 3D structure prediction of the RNP protein

To generate the model of the N protein dimer, atomic models (PDB ID: 6M3M, 6WZQ) of the N protein N-terminal domain (NTD) and C-terminal domain (CTD) were fitted into the murine hepatitis virus (MHV) N protein densities^45^ by UCSF Chimera. The theoretical model of the RNP was built in a helical manner as previously described ^45,46^.

### Prediction of the change in the polymerization state of the N protein driven by the 203 mutation

To understand the effects of the 203 mutation on the structure of the N protein, we produced three recombinant protein N variants (native, R203M and R203K) in yeast and purified them by Strep-Tactin affinity chromatography. We performed size exclusion chromatography to evaluate the polymerization state of the N variants. After injection of native N protein onto the column, two peaks were detected with retention times of 70 and 90 min. The two peaks correspond to molecular masses of ∼100 and ∼30 kDa, respectively. They are also compatible with the N protein dimer and monomer, respectively. However, the loading of R203M- or R203K-mutated protein onto the column incurred an increase in the presumed N protein dimer peak and a decrease in the presumed N protein monomer peak (Fig. S4A). Analysis of these fractions by SDS-PAGE confirmed the appearance of the N protein (Fig. S4B). Based on the predicted structure of the N protein (Fig. S4C-E), we predicted that the change in the polymerization state of the N protein driven by the 203 mutation may result in a synergistic effect on the S protein mutations, e.g., L452R.

## Supplementary information

Document S1. Figs. S1 to S5.

Table S1. Comparison of the fraction between the R203/452R and 203M/452R variants for two time points separated by a more than 2-week gap at different geographical scales.

Table S2. Comparison of the growth rates between the R203/452R and 203M/452R variants in different regional subdivisions using the Mann-Kendall trend test (MK) and isotonic regression (IR).

Table S3. Comparison of the fraction between the R203/484K/501Y and 203K/484K/501Y variants for two time points separated by a more than 2-week gap in different geographical scales.

Table S4. Comparison of the growth rates between the R203/484K/501Y and 203K/484K/501Y variants in different regional subdivisions and the trends are tested by Mann-Kendall trend test.

Table S5. Evaluation of the trends and the correlation with the vaccination coverage (VC) for the fitness of the R203M/L452R variant relative to the L452R-only variant at hierarchical geographical levels.

Table S6. Evaluation of the trends and the correlation with the vaccination coverage (VC) for the fitness of the R203M/L452R variant (Delta) relative to the R203K/N501Y variant (Alpha) at hierarchical geographical levels.

Table S7. The primers used for overlap-extension PCR in the generation of mutant viruses.

Table S8. Primers used for competition assay, viral subgenomic RNA assay and genomic RNA assay.

Table S9.Keywords used to search and annotate records with patient status.

Data S1 includes the maximum likelihood estimation in continents/countries (Secondary folder: “Maximum_Likelihood_Estimation”, legends follow Fig. 1F) and the weekly running counts of lineages in hierarchical geographic levels (Secondary folder: “Weekly_running_counts”, legends follow Fig. 1E) of the 203M/452R variant relative to the R203/452R variant (Folder: “IF_203M”) and the 203K/484K/501Y variant relative to the R203/484K/501Y variant (Folder: “IF_203K”), the trend analysis (Secondary folder: “Trend”, legends follow Fig. 4H) and correlation analysis (Secondary folder: “Cor_Vac”, legends follow Fig. 4E) results with statistical significance in hierarchical geographic levels of the fitness of the 203M/452R variant relative to the R203/452R variant (Folder: “RF_203M452R_to_203R452R”) and the 203M/452R variant relative to the 203K/501Y variant (Folder: “RF_203M452R_to_203K501Y”), the list of SARS-CoV-2 strains collected from December 2019 to September 2021 (File: “SARS2_List.xls”) and the list of clinical information of the strains collected up to September 2021 (File: “Clinical_Info.xls”).

## REFERENCES

1 Ralph, R. et al. 2019-nCoV (Wuhan virus), a novel Coronavirus: human-to-human transmission, travel-related cases, and vaccine readiness. J Infect Dev Ctries 14, 3–17, doi:10.3855/jidc.12425 (2020).

2 Lu, H., Stratton, C. W. & Tang, Y. W. Outbreak of Pneumonia of Unknown Etiology in Wuhan China: the Mystery and the Miracle. J Med Virol, doi:10.1002/jmv.25678 (2020).

3 Hui, D. S. et al. The continuing 2019-nCoV epidemic threat of novel coronaviruses to global health - The latest 2019 novel coronavirus outbreak in Wuhan, China. Int J Infect Dis 91, 264–266, doi:10.1016/j.ijid.2020.01.009 (2020).

4 Wu, F. et al. A new coronavirus associated with human respiratory disease in China. Nature 579, 265–269, doi:10.1038/s41586-020-2008-3 (2020).

5 Zhu, Z. L., Meng, K. W. & Meng, G. Genomic recombination events may reveal the evolution of coronavirus and the origin of SARS-CoV-2. Scientific reports 10, doi:10.1038/S41598-020-78703-6 (2020).

6 Smith, E. C., Blanc, H., Surdel, M. C., Vignuzzi, M. & Denison, M. R. Coronaviruses lacking exoribonuclease activity are susceptible to lethal mutagenesis: evidence for proofreading and potential therapeutics. PLoS pathogens 9, e1003565, doi:10.1371/journal.ppat.1003565 (2013).

7 Mercatelli, D. & Giorgi, F. M. Geographic and Genomic Distribution of SARS-CoV-2 Mutations. Front Microbiol 11, 1800, doi:10.3389/fmicb.2020.01800 (2020).

8 Hussain, M. et al. Structural variations in human ACE2 may influence its binding with SARS-CoV-2 spike protein. Journal of medical virology 92, 1580–1586, doi:10.1002/jmv.25832 (2020).

9 Salvatori, G. et al. SARS-CoV-2 SPIKE PROTEIN: an optimal immunological target for vaccines. Journal of translational medicine 18, 222, doi:10.1186/s12967-020-02392-y (2020).

10 Korber, B. et al. Tracking Changes in SARS-CoV-2 Spike: Evidence that D614G Increases Infectivity of the COVID-19 Virus. Cell 182, 812–827 e819, doi:10.1016/j.cell.2020.06.043 (2020).

11 Plante, J. A. et al. Spike mutation D614G alters SARS-CoV-2 fitness. Nature, doi:10.1038/s41586-020-2895-3 (2020).

12 Zhu, Z. et al. Rapid Spread of Mutant Alleles in Worldwide SARS-CoV-2 Strains Revealed by Genome-Wide Single Nucleotide Polymorphism and Variation Analysis. Genome biology and evolution 13, doi:10.1093/gbe/evab015 (2021).

13 Mok, B. W.-Y. et al. SARS-CoV-2 spike D614G variant exhibits highly efficient replication and transmission in hamsters. bioRxiv (2020).

14 Trucchi, E. et al. Population Dynamics and Structural Effects at Short and Long Range Support the Hypothesis of the Selective Advantage of the G614 SARS-CoV-2 Spike Variant. Mol Biol Evol 38, 1966–1979, doi:10.1093/molbev/msaa337 (2021).

15 Cheng, L. et al. Impact of the N501Y substitution of SARS-CoV-2 Spike on neutralizing monoclonal antibodies targeting diverse epitopes. Virology journal 18, 87, doi:10.1186/s12985-021-01554-8 (2021).

16 Zhao, S. et al. Quantifying the transmission advantage associated with N501Y substitution of SARS-CoV-2 in the United Kingdom: An early data-driven analysis. Journal of travel medicine, doi:10.1093/jtm/taab011 (2021).

17 Garcia-Beltran, W. F. et al. Multiple SARS-CoV-2 variants escape neutralization by vaccine-induced humoral immunity. Cell, doi:10.1016/j.cell.2021.03.013 (2021).

18 Liu, Y. et al. The N501Y spike substitution enhances SARS-CoV-2 infection and transmission. Nature, doi:10.1038/s41586-021-04245-0 (2021).

19 Jangra, S. et al. SARS-CoV-2 spike E484K mutation reduces antibody neutralisation. The Lancet. Microbe, doi:10.1016/S2666-5247(21)00068-9 (2021).

20 McCallum, M. et al. SARS-CoV-2 immune evasion by variant B.1.427/B.1.429. bioRxiv, doi:10.1101/2021.03.31.437925 (2021).

21 Motozono, C. et al. SARS-CoV-2 spike L452R variant evades cellular immunity and increases infectivity. Cell host & microbe 29, 1124–1136 e1111, doi:10.1016/j.chom.2021.06.006 (2021).

22 Deng, X. et al. Transmission, infectivity, and neutralization of a spike L452R SARS-CoV-2 variant. Cell 184, 3426–3437 e3428, doi:10.1016/j.cell.2021.04.025 (2021).

23 Wu, H. et al. Nucleocapsid mutations R203K/G204R increase the infectivity, fitness, and virulence of SARS-CoV-2. Cell host & microbe, doi:10.1016/j.chom.2021.11.005 (2021).

24 Rochman, N. D. et al. Ongoing global and regional adaptive evolution of SARS-CoV-2. Proc Natl Acad Sci U S A 118, doi:10.1073/pnas.2104241118 (2021).

25 Syed, A. M. et al. Rapid assessment of SARS-CoV-2 evolved variants using virus-like particles. Science, eabl6184, doi:10.1126/science.abl6184 (2021).

26 Mlcochova, P. et al. SARS-CoV-2 B.1.617.2 Delta variant replication and immune evasion. Nature 599, 114–119, doi:10.1038/s41586-021-03944-y (2021).

27 West, A. P., Jr. et al. Detection and characterization of the SARS-CoV-2 lineage B.1.526 in New York. Nature communications 12, 4886, doi:10.1038/s41467-021-25168-4 (2021).

28 Wang, Y. et al. New framework for recombination and adaptive evolution analysis with application to the novel coronavirus SARS-CoV-2. Briefings in bioinformatics 22, doi:10.1093/bib/bbab107 (2021).

29 Mishra, A. et al. SARS-CoV-2 Delta Variant among Asiatic Lions, India. Emerg Infect Dis 27, 2723–2725, doi:10.3201/eid2710.211500 (2021).

30 Goldberg, Y. et al. Waning Immunity after the BNT162b2 Vaccine in Israel. The New England journal of medicine 385, e85, doi:10.1056/NEJMoa2114228 (2021).

31 Saito, A. et al. Enhanced fusogenicity and pathogenicity of SARS-CoV-2 Delta P681R mutation. Nature, doi:10.1038/s41586-021-04266-9 (2021).

32 Petersen, E. et al. Emergence of new SARS-CoV-2 Variant of Concern Omicron (B.1.1.529) - highlights Africa’s research capabilities, but exposes major knowledge gaps, inequities of vaccine distribution, inadequacies in global COVID-19 response and control efforts. International journal of infectious diseases : IJID : official publication of the International Society for Infectious Diseases, doi:10.1016/j.ijid.2021.11.040 (2021).

33 Kumar, S., Thambiraja, T. S., Karuppanan, K. & Subramaniam, G. Omicron and Delta Variant of SARS-CoV-2: A Comparative Computational Study of Spike protein. bioRxiv (2021).

34 Zhu, Z., Meng, K., Liu, G. & Meng, G. A database resource and online analysis tools for coronaviruses on a historical and global scale. Database : the journal of biological databases and curation, doi:10.1093/database/baaa070 (2020).

35 Volz, E. et al. Evaluating the Effects of SARS-CoV-2 Spike Mutation D614G on Transmissibility and Pathogenicity. Cell 184, 64–75 e11, doi:10.1016/j.cell.2020.11.020 (2021).

36 Suchard, M. A. et al. Bayesian phylogenetic and phylodynamic data integration using BEAST 1.10. Virus evolution 4, vey016, doi:10.1093/ve/vey016 (2018).

37 Price, M. N., Dehal, P. S. & Arkin, A. P. FastTree 2--approximately maximum-likelihood trees for large alignments. PLoS One 5, e9490, doi:10.1371/journal.pone.0009490 (2010).

38 Volz, E. M. & Didelot, X. Modeling the Growth and Decline of Pathogen Effective Population Size Provides Insight into Epidemic Dynamics and Drivers of Antimicrobial Resistance. Syst Biol 67, 719–728, doi:10.1093/sysbio/syy007 (2018).

39 Xie, X. et al. An Infectious cDNA Clone of SARS-CoV-2. Cell host & microbe 27, 841–848 e843, doi:10.1016/j.chom.2020.04.004 (2020).

40 Xie, X. et al. Engineering SARS-CoV-2 using a reverse genetic system. Nature protocols 16, 1761–1784, doi:10.1038/s41596-021-00491-8 (2021).

41 Corman, V. M. et al. Detection of 2019 novel coronavirus (2019-nCoV) by real-time RT-PCR. Euro surveillance : bulletin Europeen sur les maladies transmissibles = European communicable disease bulletin 25, doi:10.2807/1560-7917.ES.2020.25.3.2000045 (2020).

42 Dagotto, G. et al. Comparison of Subgenomic and Total RNA in SARS-CoV-2 Challenged Rhesus Macaques. Journal of virology, doi:10.1128/JVI.02370-20 (2021).

43 Hamilton, M. B. Population Genetics. (A John Wiley & Sons, Ltd., Publication, 2009).

44 Zhang, Y. et al. A gRNA-tRNA array for CRISPR-Cas9 based rapid multiplexed genome editing in Saccharomyces cerevisiae. Nat. Commun. 10, 1053, doi:10.1038/s41467-019-09005-3 (2019).

45 Gui, M. et al. Electron microscopy studies of the coronavirus ribonucleoprotein complex. Protein Cell 8, 219–224, doi:10.1007/s13238-016-0352-8 (2017).

46 Wu, H. et al. Nucleocapsid mutation R203K/G204R increases the infectivity, fitness and virulence of SARS-CoV-2. bioRxiv, 2021.2005.2024.445386, doi:10.1101/2021.05.24.445386 (2021).

