## Supplementary Figs. S1-S5 for "Nucleocapsid 203 mutations enhance SARS-CoV-2 immune evasion"

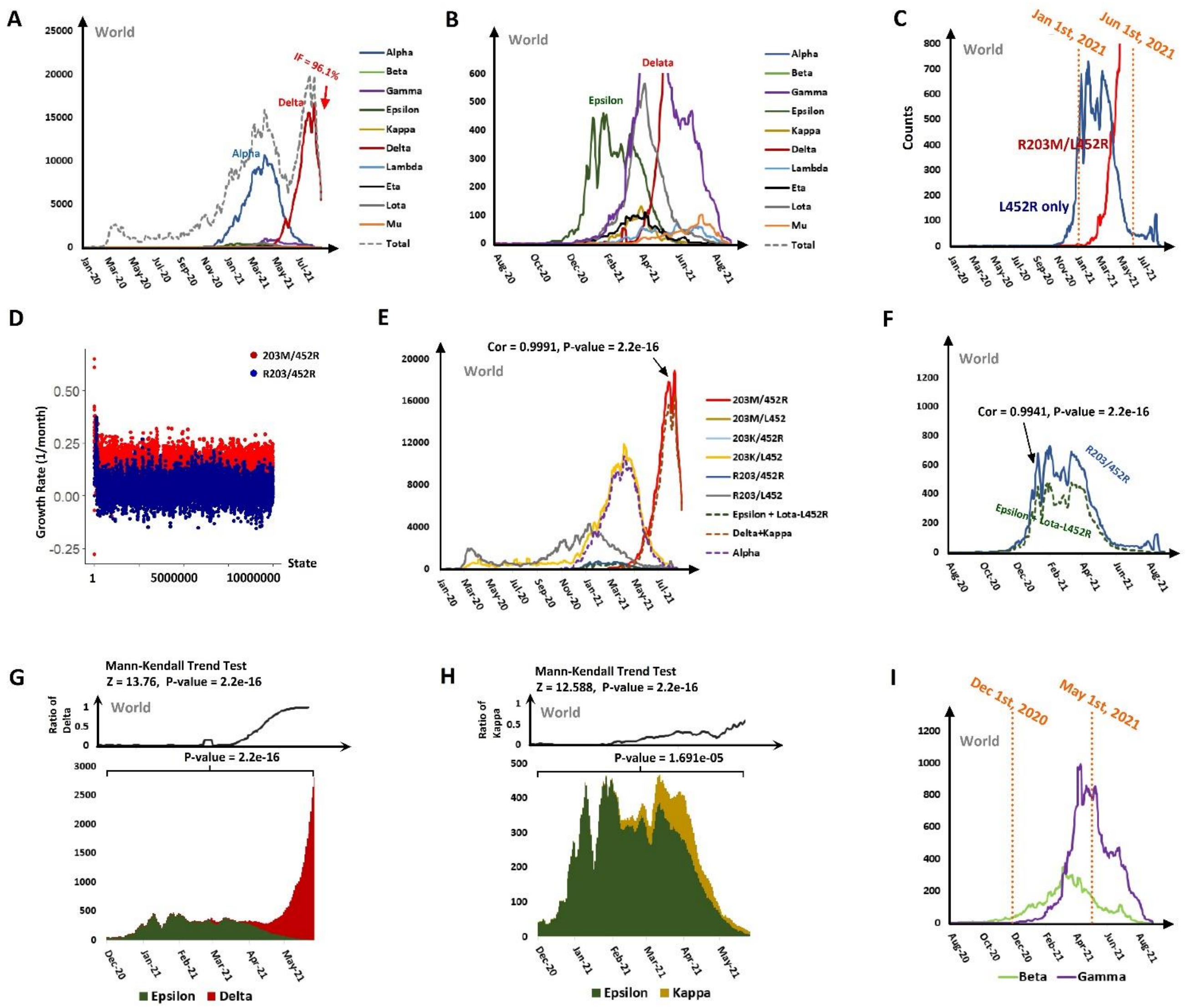


Fig. S1. Comparison of the S protein mutation variants with the nucleotide 203 mutation and without the 203 mutation, corresponded to Fig. 1. (A) Weekly running counts of lineages of concern from December 2019 to September 2021. (B) An enlarged image showing the weekly running counts of the lineages that used to have a rapid spread but did not achieve a dominance around the world. (C) Comparison of the weekly running counts between the R203M/L452R and L452R-only variants with the temporal vicinity of the introduction of the 203M/452R variant (January 1st, 2021) and the date six months later (June 1st, 2021) marked. (D) The growth rates of the strains across states in an exponential growth model. (E) Worldwide weekly running counts of the 6 combinations of the N protein 203 mutations and the S protein 452 mutations, as well as those of the Epsilon+Lota-L452R, Delta+Kappa and Alpha variants. The correlation between the counts of the 203M/452R and Delta+Kappa variants is shown at the top. (F) Worldwide weekly running counts of the R203/452R and Epsilon+Lota-L452R variants. The correlation between them was calculated and is presented at the top. The upper panel shows the changes in the ratio of Delta in Delta and Epsilon with the result of the Mann-Kendall trend test listed at the top. The lower panel shows the weekly running counts of Delta and Epsilon from December 2020 to June 2021. The result of Fisher's exact test at the time interval bracketed is shown at the top. The analysis in (H) is similar to that in (G), but the targets are Epsilon and Kappa. (I) Comparison of the weekly running counts between Beta (R203/484K/501Y) and Gamma (203K/484K/501Y) with the approximately initial global rise of Gamma (December 1st, 2020) and the date six months later (May 1st, 2021) marked.

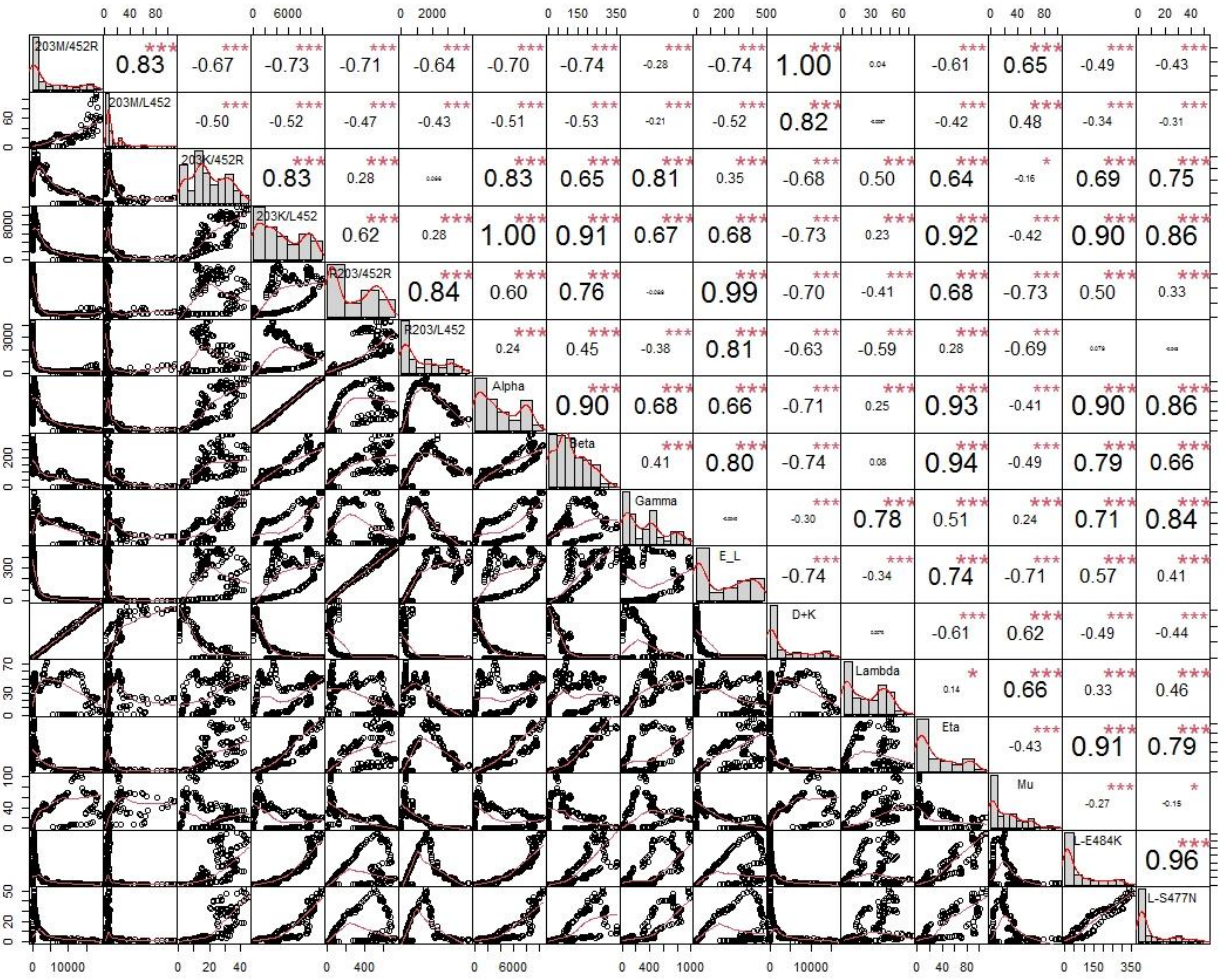


Fig. S2. Correlation tests between pairs of lineages, corresponded to Fig. 1.

The items in comparison include the 6 combinations of the N protein 203 mutation and the S protein 452 mutation and the main VOCs. E_L denotes the lineages Epsilon+Lota-452R. D+K denotes the lineages Delta+Kappa. L-E484K and L-S477N denote Lota-E484K and Lota-S477N, respectively. “*”, “**” and “***” denote P-value <0.1, P-value <0.05 and P-value < 0.01, respectively.

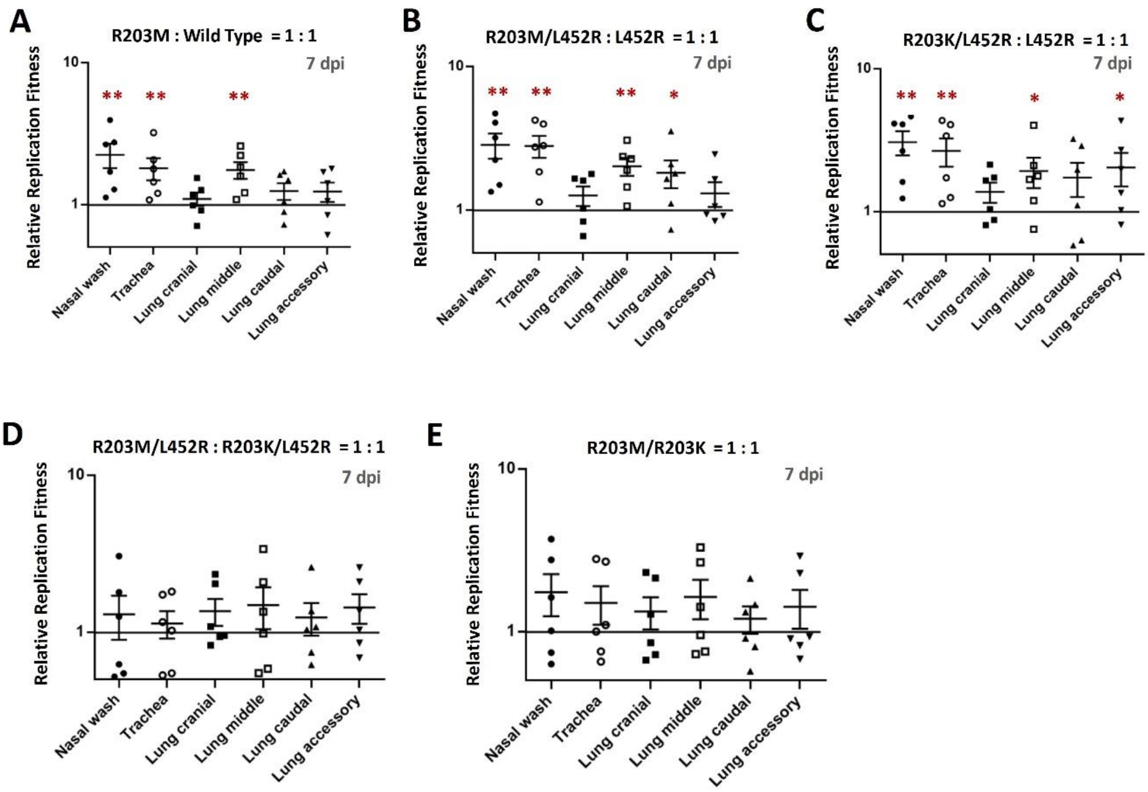


Fig. S3 Competition experiments in hamsters at 7 dpi, corresponded to Fig. 2.

(A-E) Hamsters were intranasally infected with a 1:1 mixture of wild type or mutant viruses as indicated. Nasal washes, tracheas and lungs were collected on day 7 after infection. The relative RNA amounts of R203M : wild type (A), L452R/R203M : L452R (B), L452R/R203K : L452R (C), L452R/R203M : L452R/R203K (D) and R203M : R203K (E) were assessed by RT-PCR and Sanger sequencing. Log10 scale was used for the Y-axis. Data are presented as the mean ± s.e.m.. Dots represent individual hamsters (n = 6). *, p<0.05, **, p<0.01.

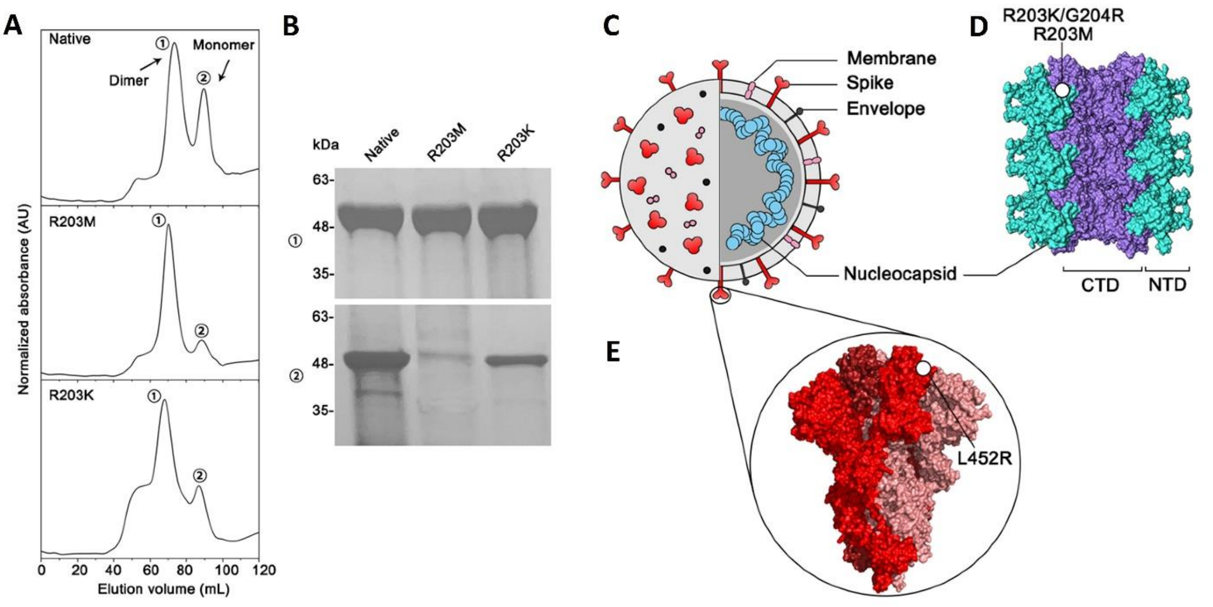


Fig. S4 Evidence showing the change in polymerization state in the nucleocapsid.

(A, B) Size exclusion chromatography of N protein variants. The elution profile of N protein was obtained by size exclusion chromatography and analysed by SDS-PAGE. (A) Injection of N protein resulted in two peaks corresponding to the dimer and monomer. The R203M and R203K mutations result in an increase in the proportion of N protein dimers. (B) The fraction containing N protein was collected, concentrated, and subjected to SDS-PAGE, which showed prominent bands at 48 kDa. (C-E) Structural representation of the mutations in the S protein and the N protein. (C) Structure of SARS-Cov-2 depicting the structural proteins. (D) A RNP protein fragment model, with the CTD depicted in purple, and the NTD in cyan. The circle indicates the R203K/G204R or R203M mutation site. Here, only one of the mutation sites is shown. (E) The spike complex is shown in the "all-down" conformation. One of its L452R mutation sites is depicted by a circle. Images in (D)-(E) were prepared using Chimera.

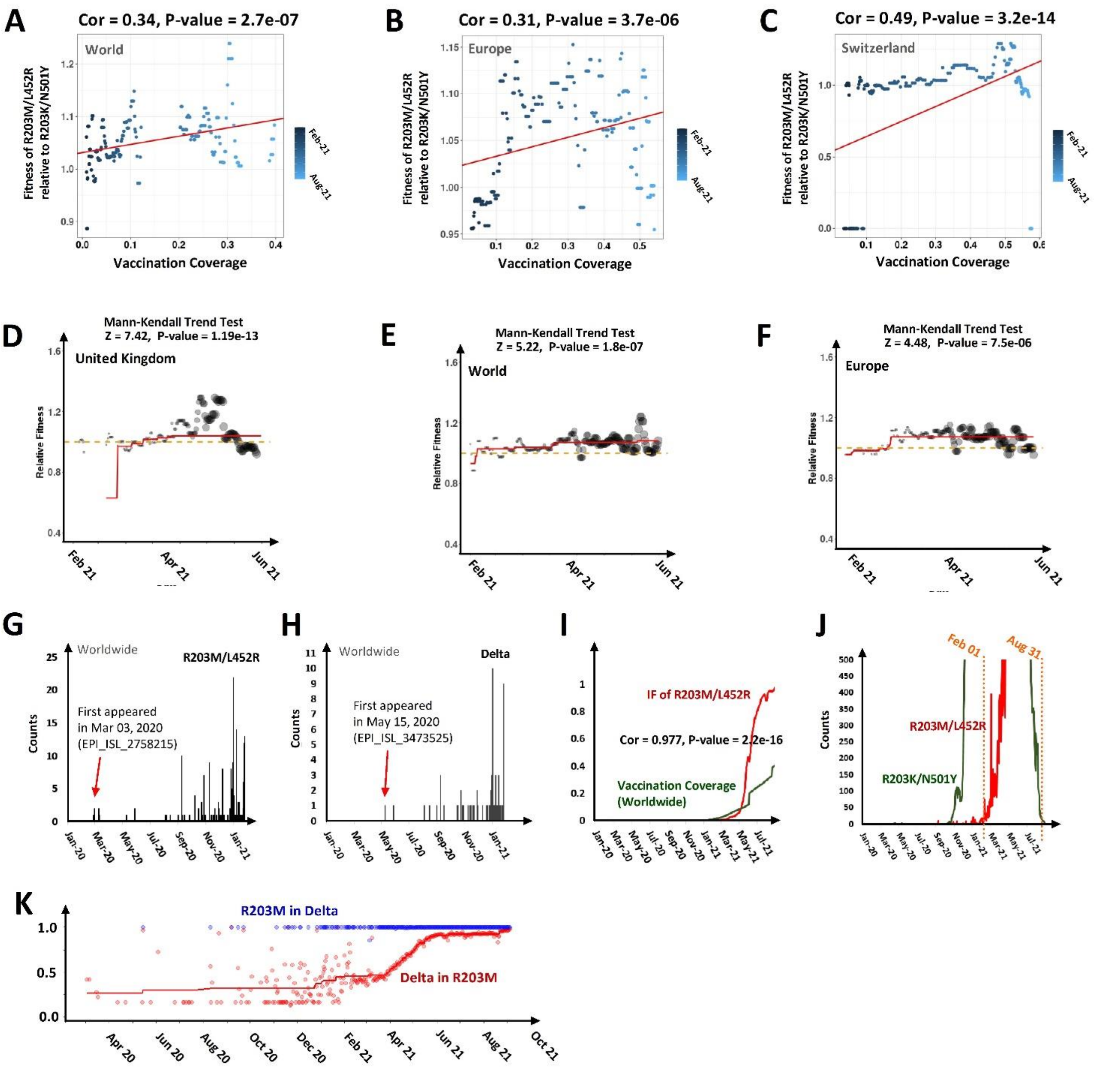


Fig. S5 Increased fitness of the R203M/L452R variant relative to that of the R203K/N501Y variant, corresponded to Fig. 4.

(A-C) Correlation analysis results between the fitness of the R203M/L452R variant relative to that of the R203K/N501Y variant (y-axis) and VC (x-axis) in the world (A), in Europe (B) and in Switzerland (C). (D-F) The fitted trend of the fitness of the R203M/L452R variant relative to that of the R203K/N501Y variant worldwide (D), in Europe (E) and in Switzerland (F). The result of the Mann-Kendall trend test is listed at the top. (G) and (H) are the counts of the R203M/L452R variant and Delta before the expansion in 2021, respectively. The first identified strain is marked by a red arrow. (I) The figure contemporarily shows the IF of the R203M/L452R variant and worldwide VC. The correlation between them was 0.977 with statistical significance. (J) Comparison of the weekly running counts between the R203M/L452R and R203K/N501Y variants with the time interval for the relative fitness calculation marked by orange dotted lines. (K) The fractions of Delta in the R203M variants (red) and the fractions of R203M in the Delta variants (blue). The calculation is based on weekly running counts.
