## Supplementary figures and images for "Nucleocapsid 203 mutations enhance SARS-CoV-2 immune evasion"

### Africa.jpeg

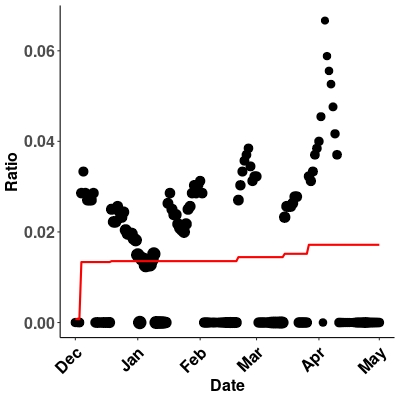

### Africa.jpeg

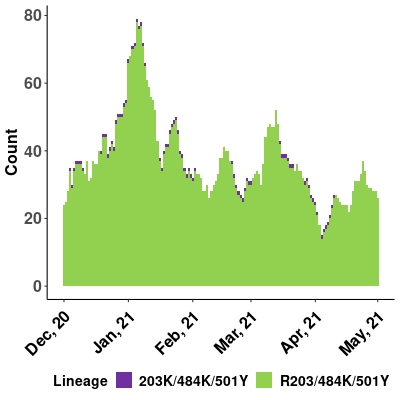

### Africa.jpeg

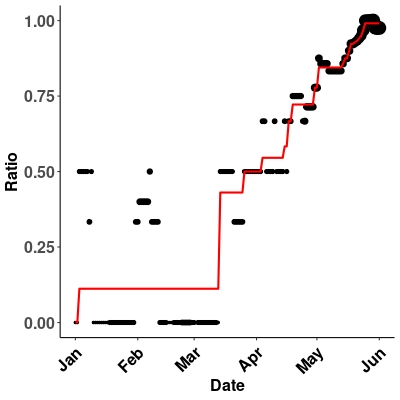

### Asia.jpeg

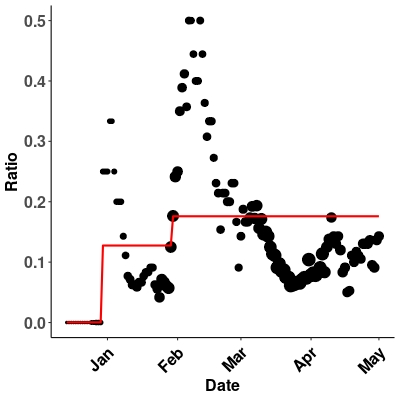

### Asia.jpeg

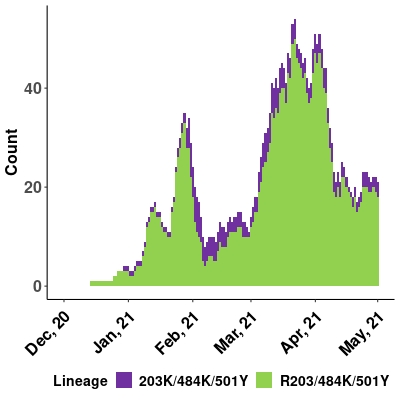

### Asia.jpeg

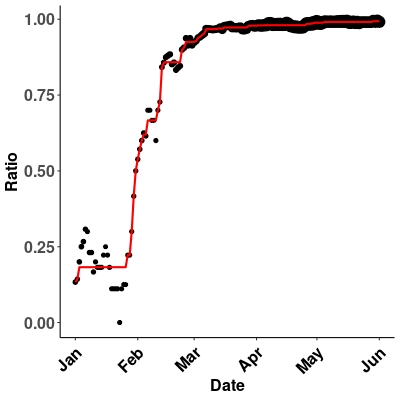

### Australia.jpeg

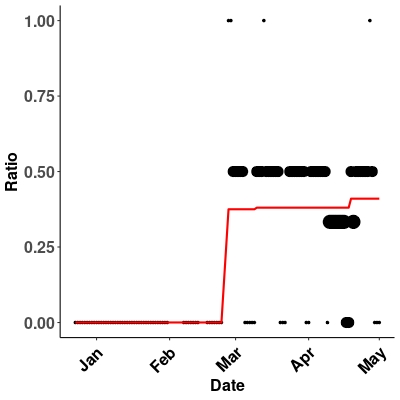

### Australia.jpeg

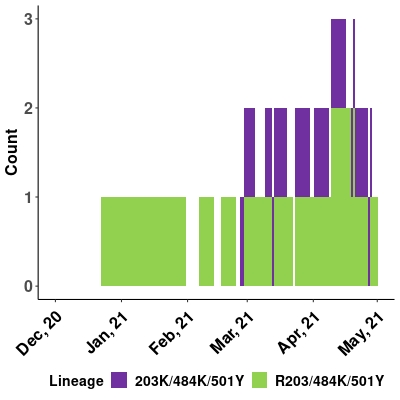

### Australia.jpeg

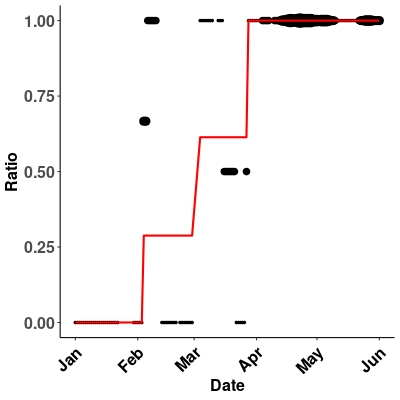

### Belgium.jpeg

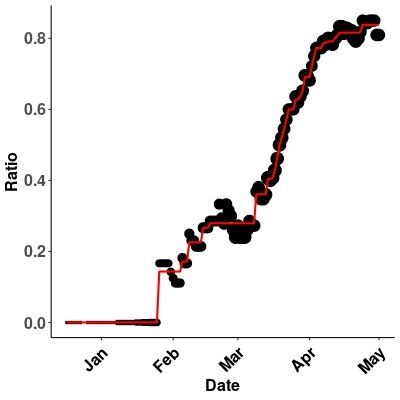

### Belgium.jpeg

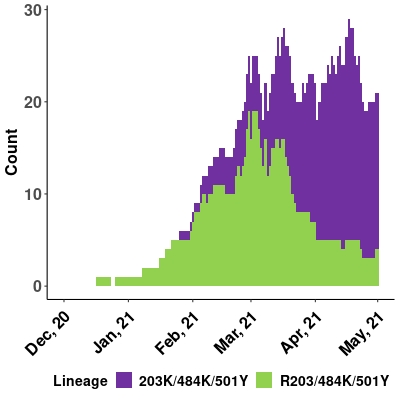

### Belgium.jpeg

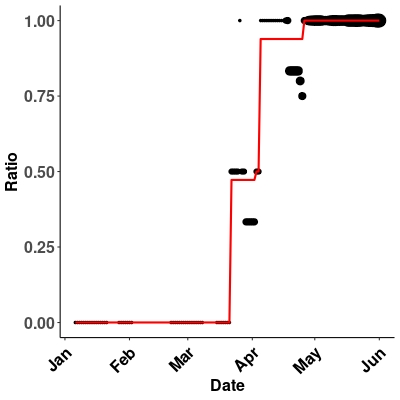

### Canada.jpeg

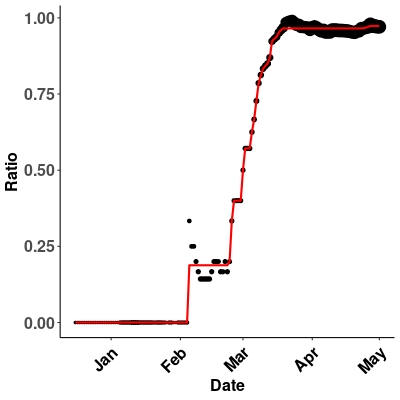

### Canada.jpeg

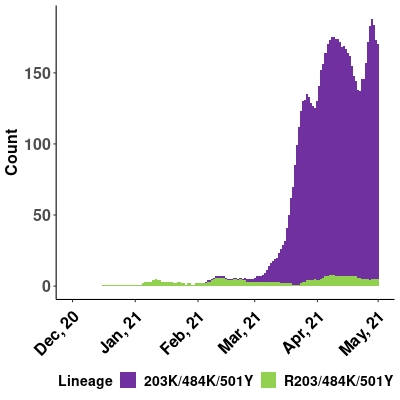

### Canada.jpeg

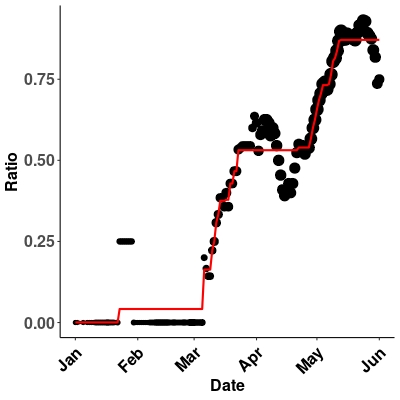

### Costa_Rica.jpeg

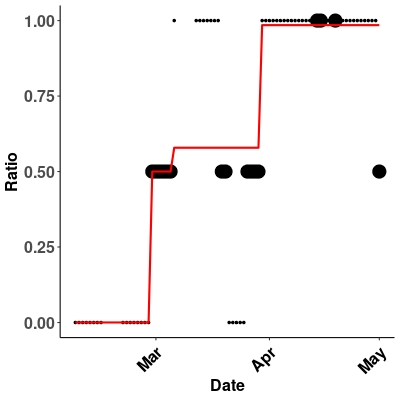

### Costa_Rica.jpeg

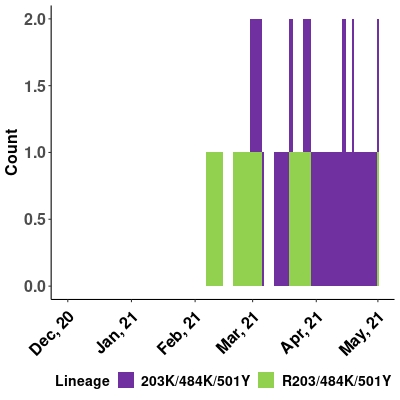

### Denmark.jpeg

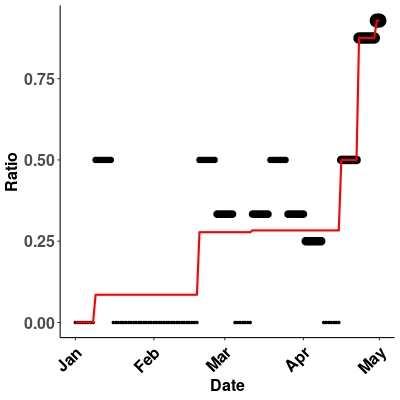

### Denmark.jpeg

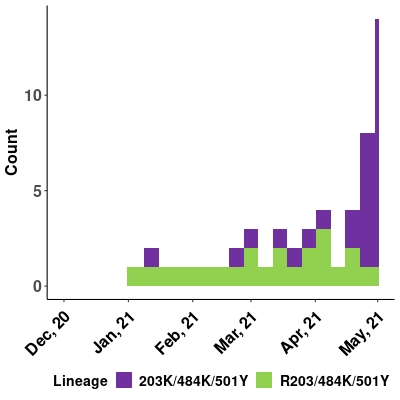

### Denmark.jpeg

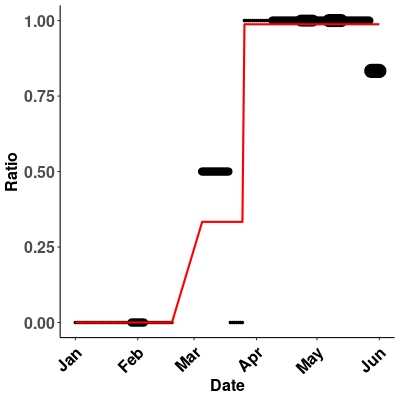

### Europe.jpeg

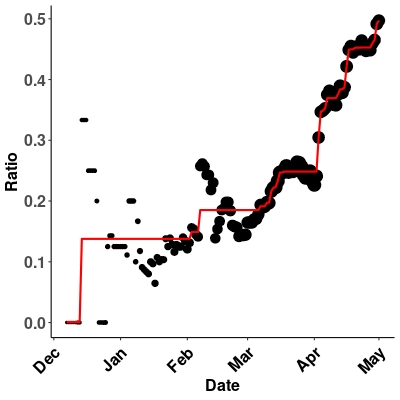

### Europe.jpeg

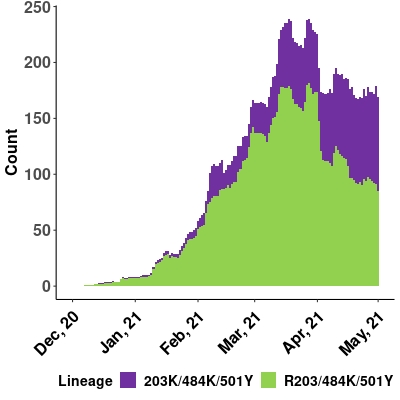

### Europe.jpeg

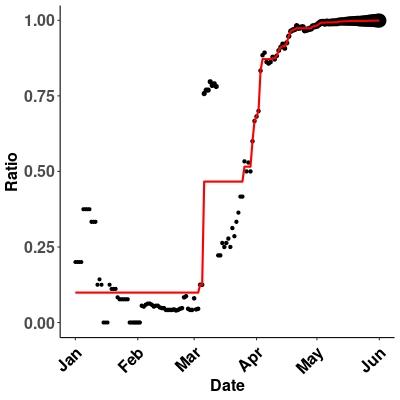

### Finland.jpeg

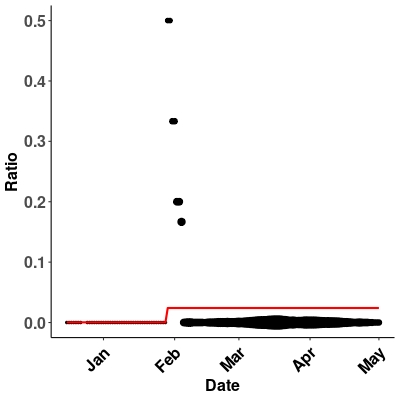

### Finland.jpeg

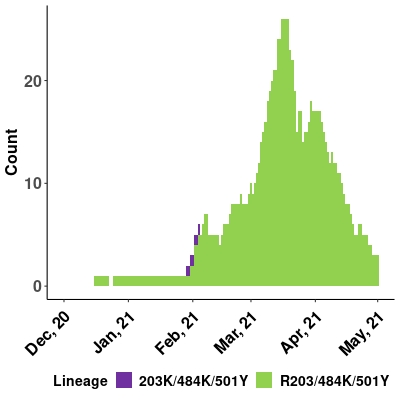

### Finland.jpeg

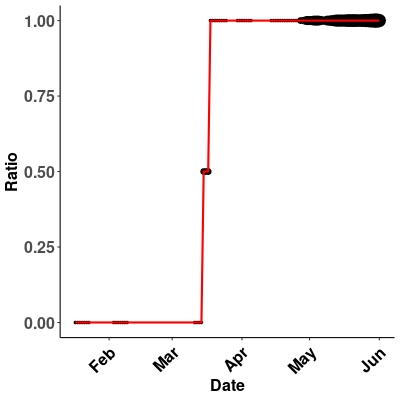

### France.jpeg

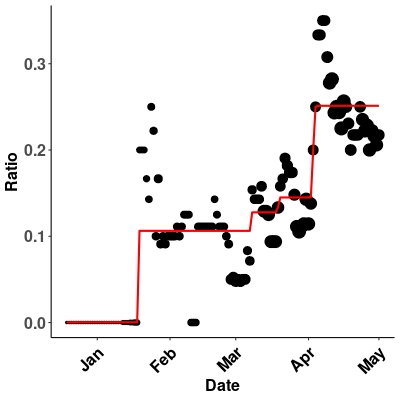

### France.jpeg

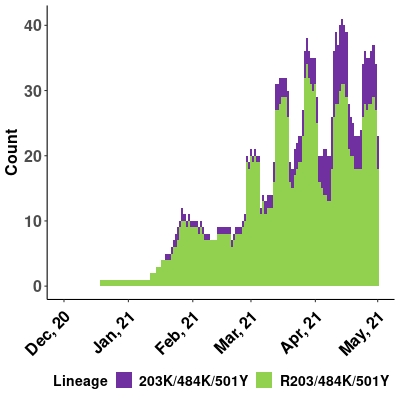

### France.jpeg

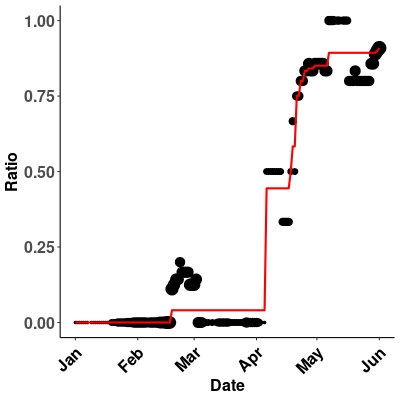

### Germany.jpeg

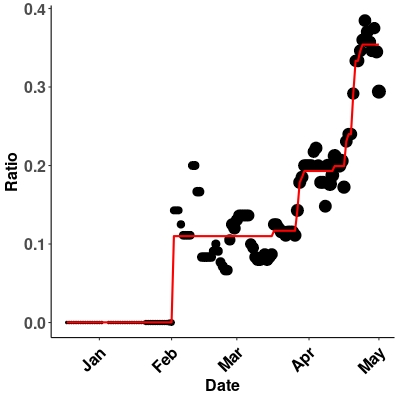

### Germany.jpeg

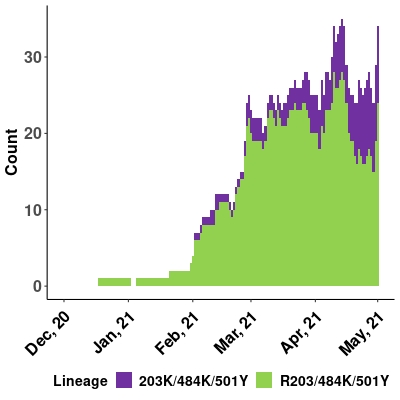

### Germany.jpeg

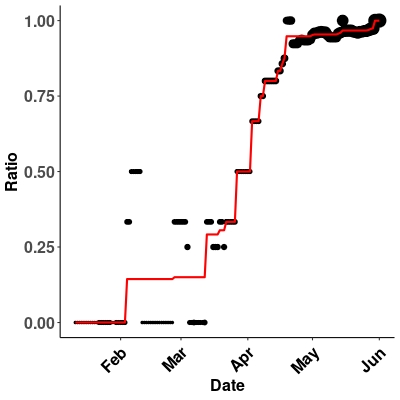

### India.jpeg

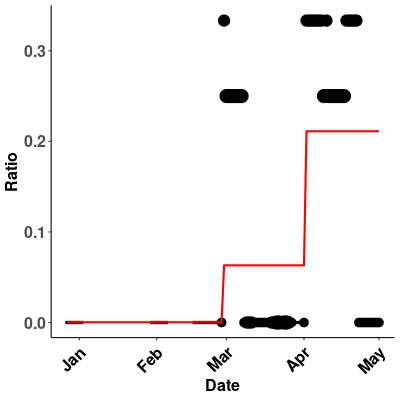

### India.jpeg

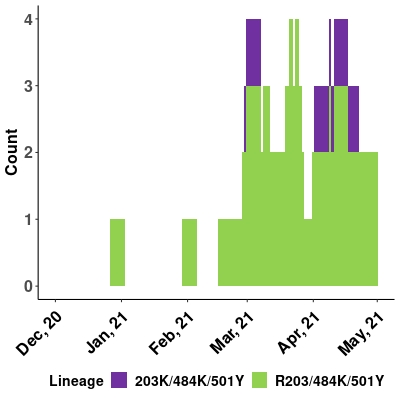

### India.jpeg

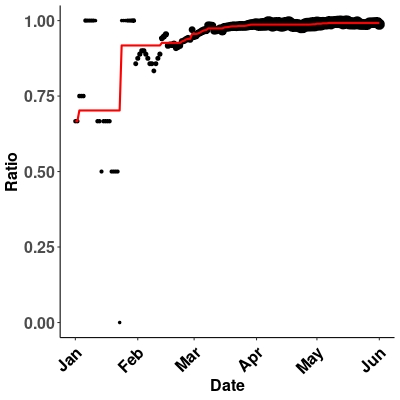

### Indonesia.jpeg

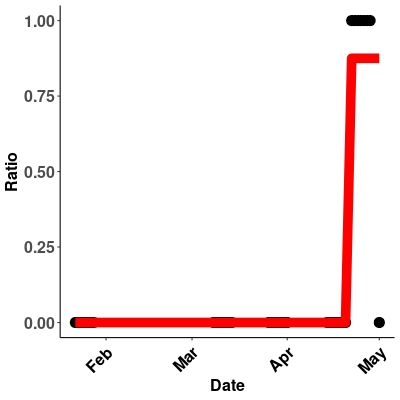

### Indonesia.jpeg

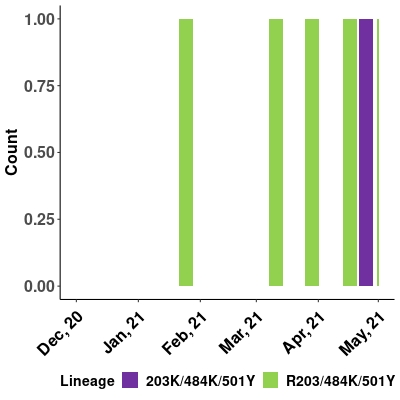

### Indonesia.jpeg

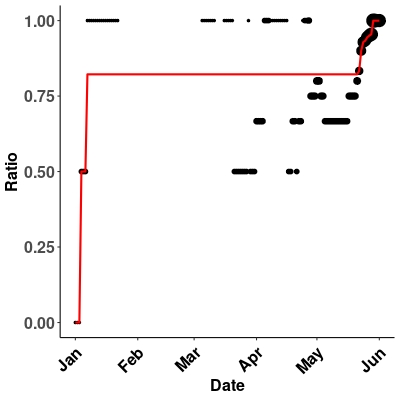

### Israel.jpeg

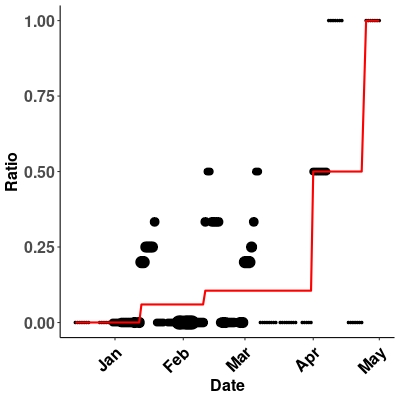

### Israel.jpeg

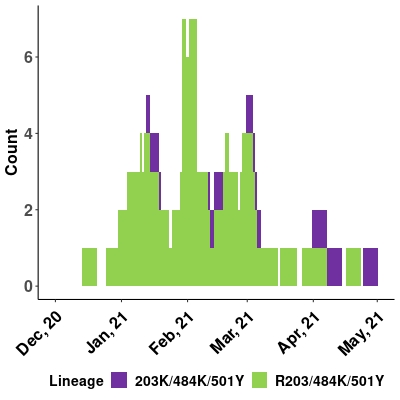

### Israel.jpeg

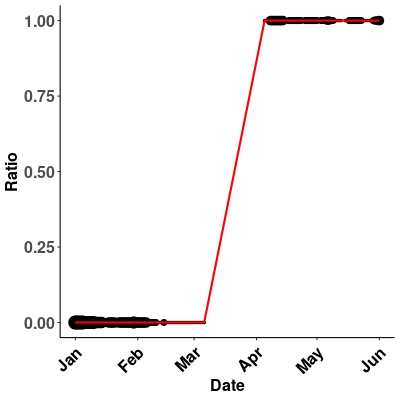

### Italy.jpeg

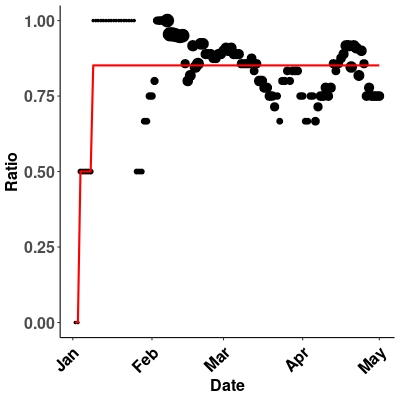

### Italy.jpeg

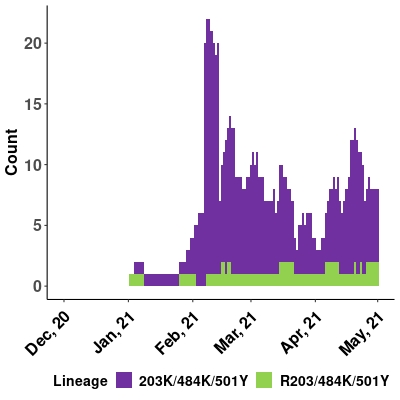

### Italy.jpeg

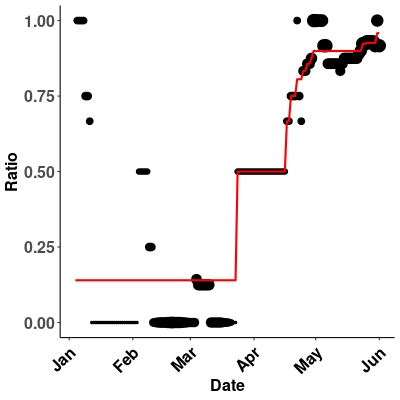

### Japan.jpeg

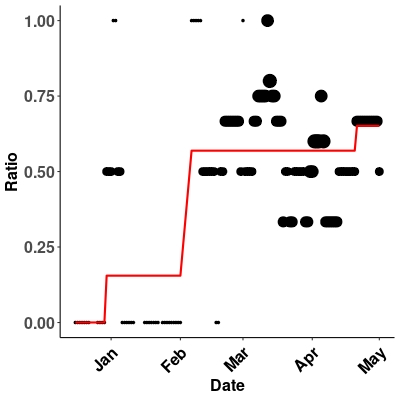

### Japan.jpeg

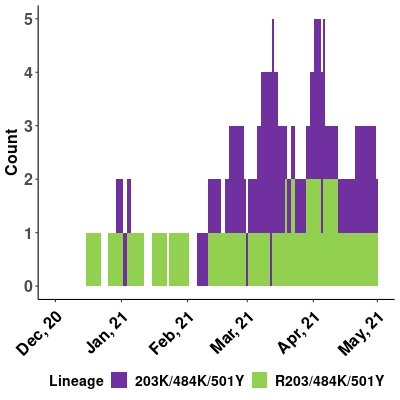

### Japan.jpeg

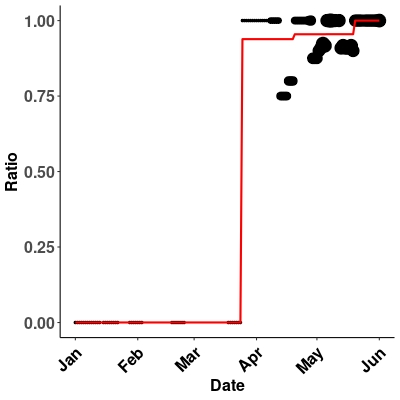

### Kenya.jpeg

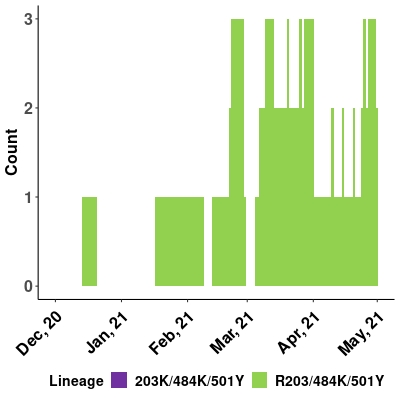

### Kenya.jpeg

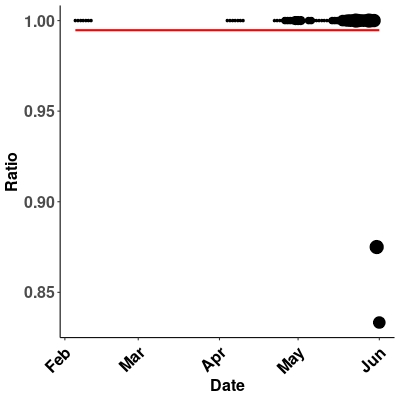

### Latin_America.jpeg

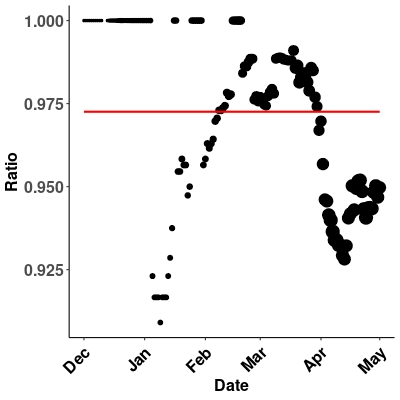

### Latin_America.jpeg

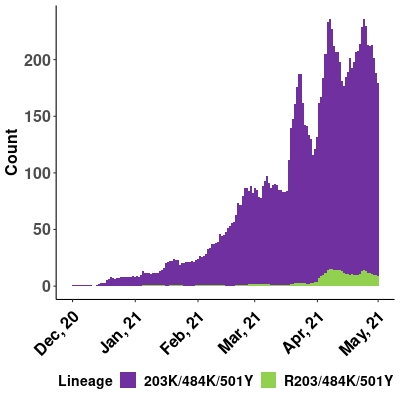

### Latin_America.jpeg

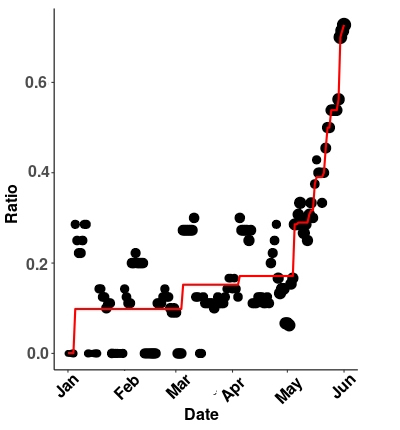

### Luxembourg.jpeg

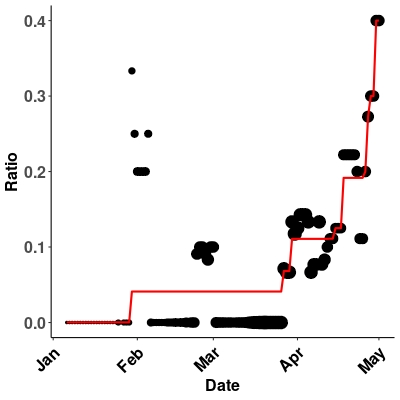

### Luxembourg.jpeg

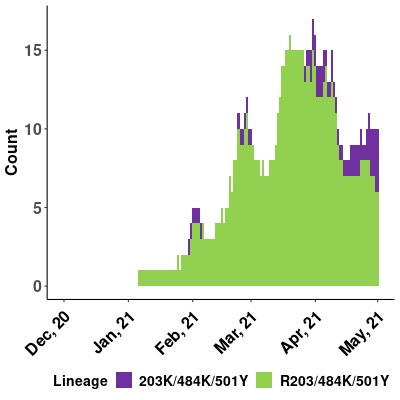

### Luxembourg.jpeg

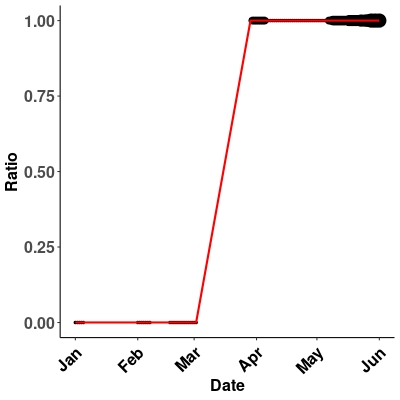

### Mexico.jpeg

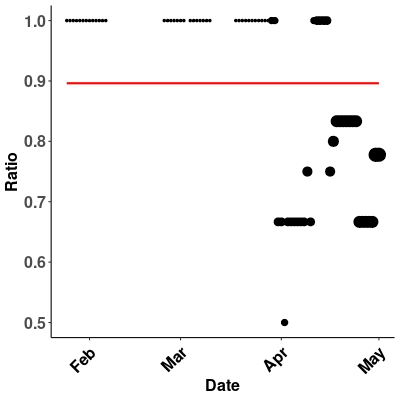

### Mexico.jpeg

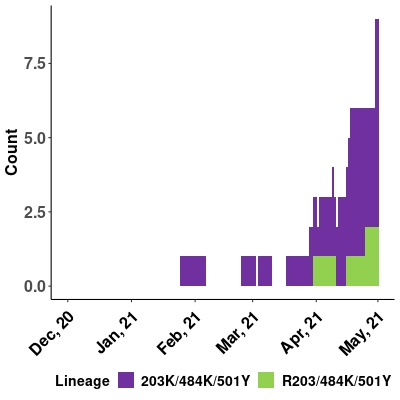

### Mexico.jpeg

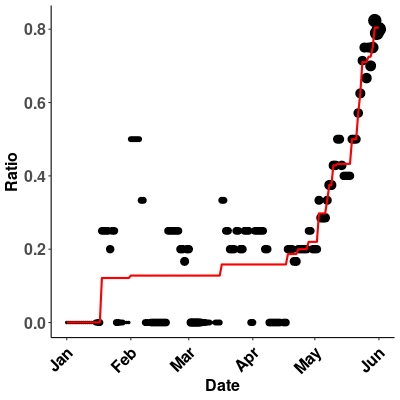

### Netherlands.jpeg

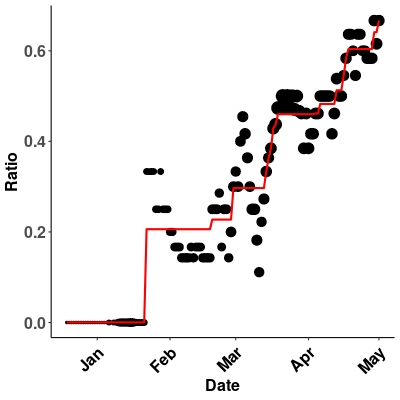

### Netherlands.jpeg

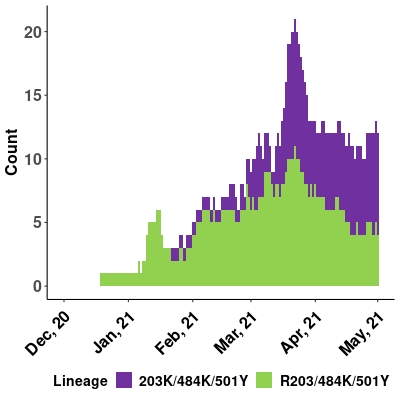

### Netherlands.jpeg

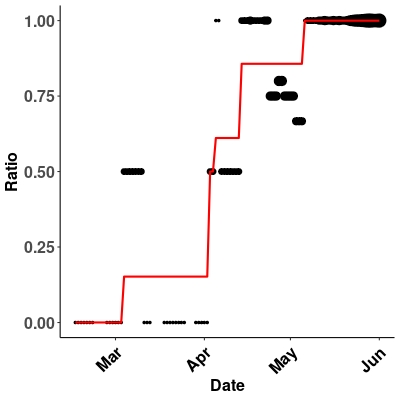

### North_America.jpeg

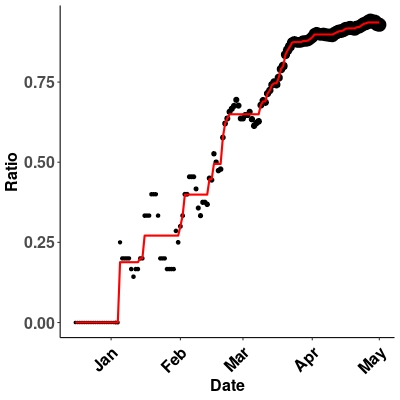

### North_America.jpeg

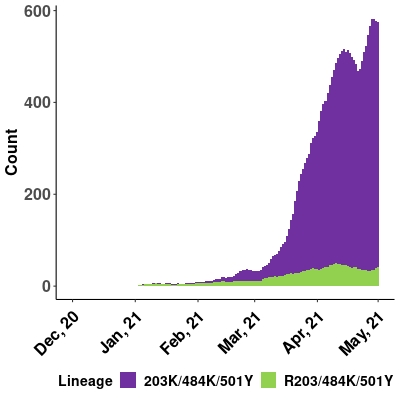

### North_America.jpeg

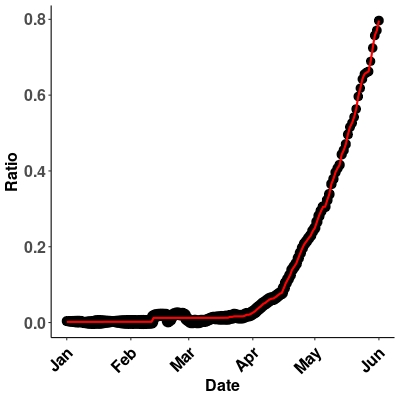

### Norway.jpeg

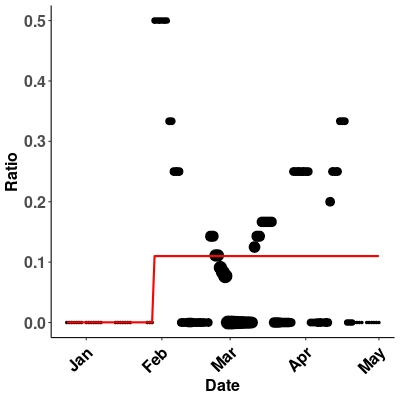

### Norway.jpeg

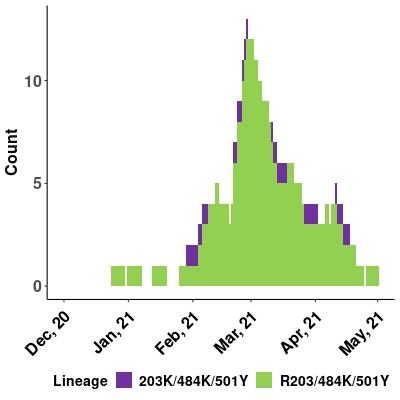

### Norway.jpeg

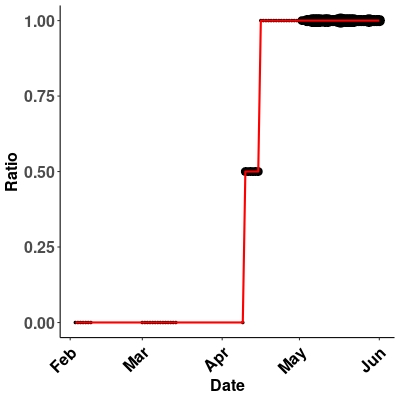

### Oceania.jpeg

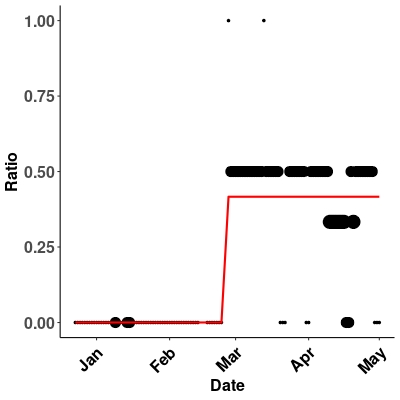

### Oceania.jpeg

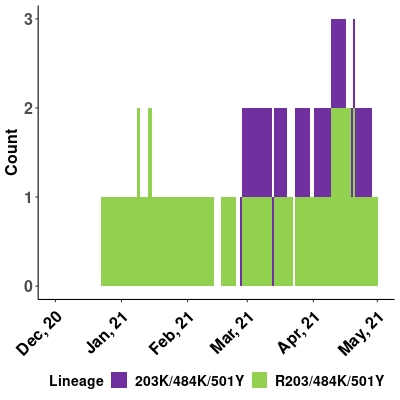

### Oceania.jpeg

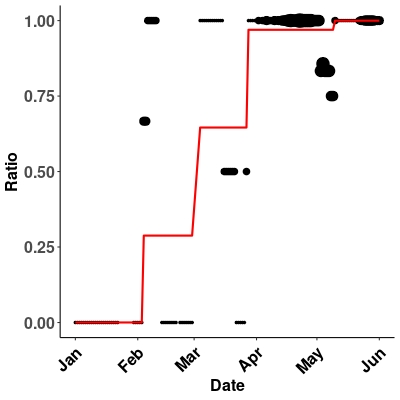

### Panama.jpeg

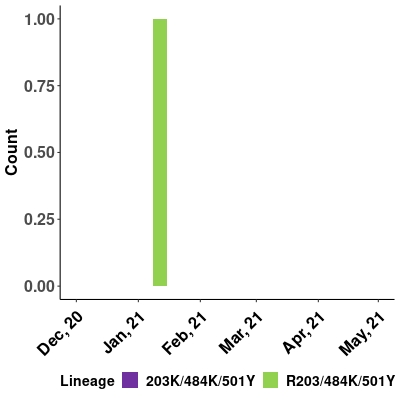

### Portugal.jpeg

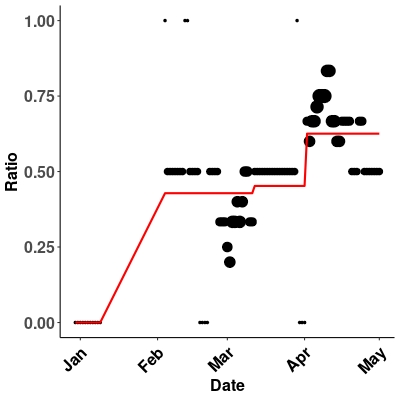

### Portugal.jpeg

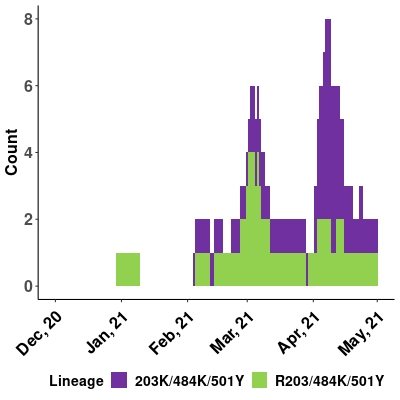

### Portugal.jpeg

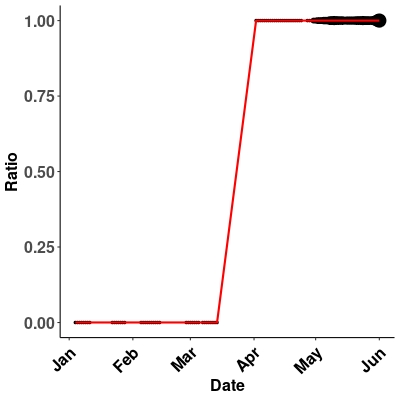

### Qatar.jpeg

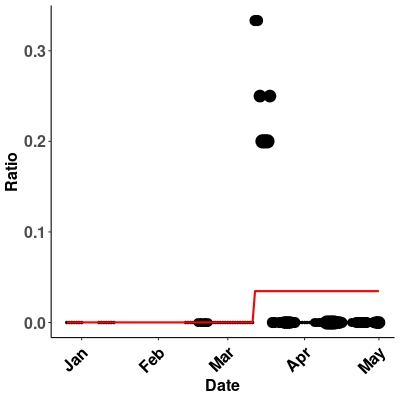

### Qatar.jpeg

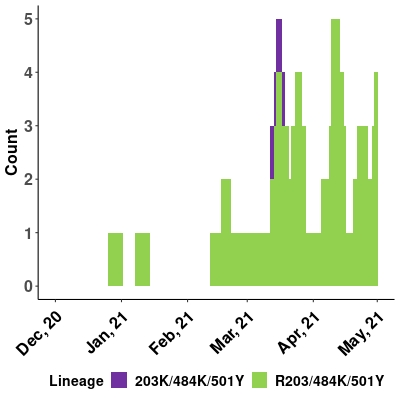

### Qatar.jpeg

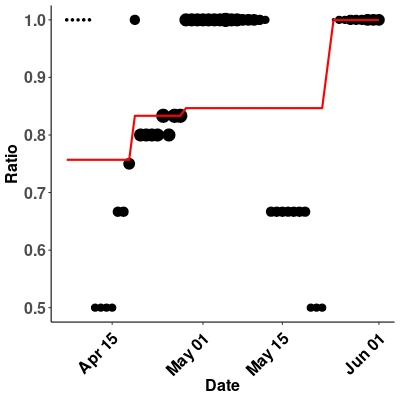

### Russia.jpeg

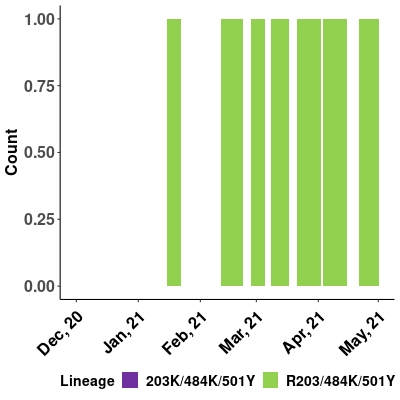

### Russia.jpeg

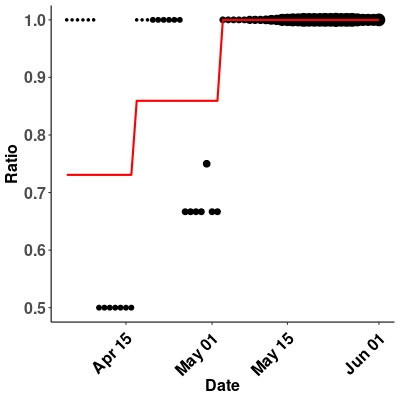

### Singapore.jpeg

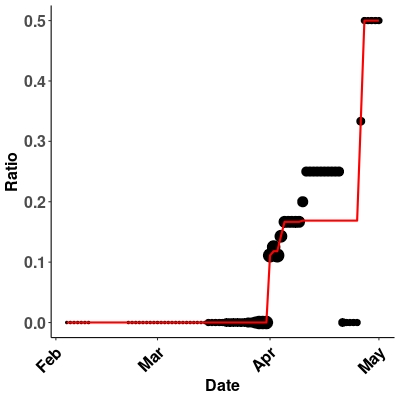

### Singapore.jpeg

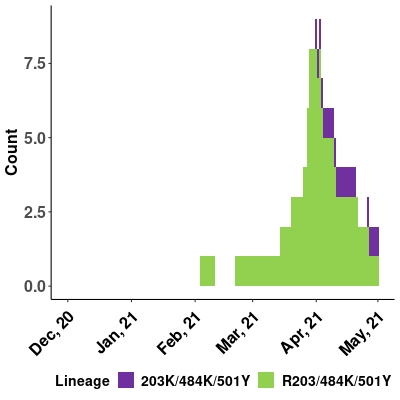

### Singapore.jpeg

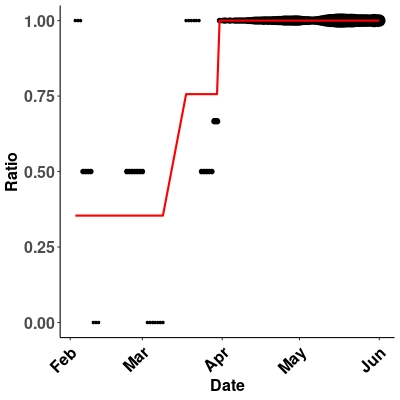

### South_Africa.jpeg

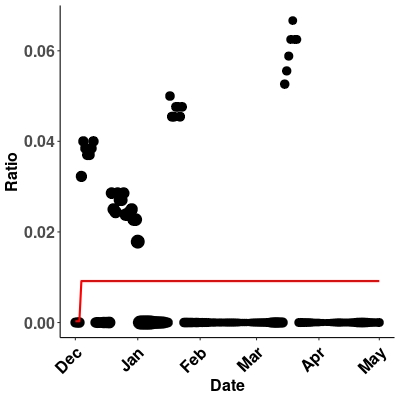

### South_Africa.jpeg

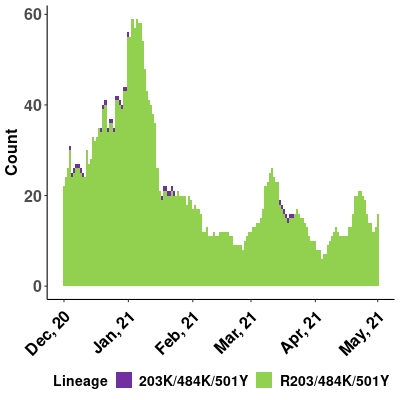

### South_Korea.jpeg

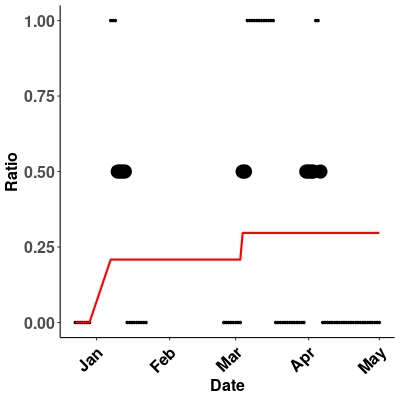

### South_Korea.jpeg

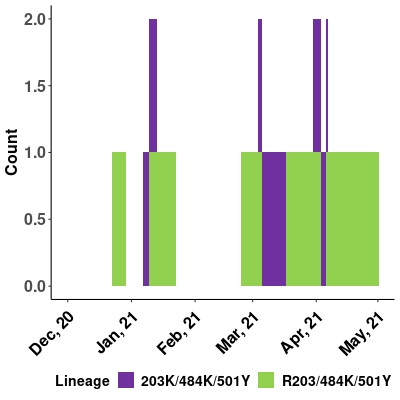

### Spain.jpeg

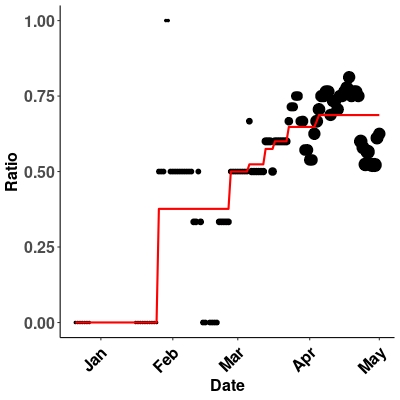

### Spain.jpeg

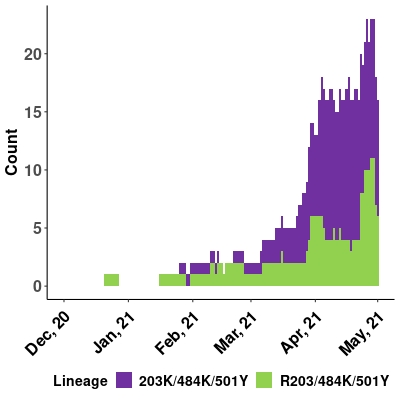

### Sweden.jpeg

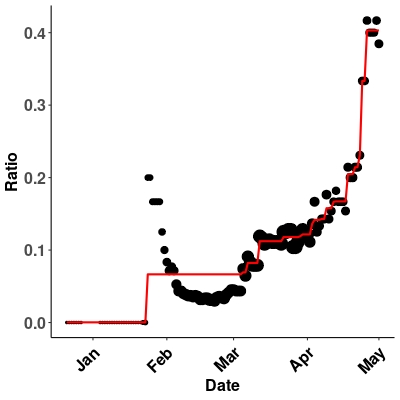

### Sweden.jpeg

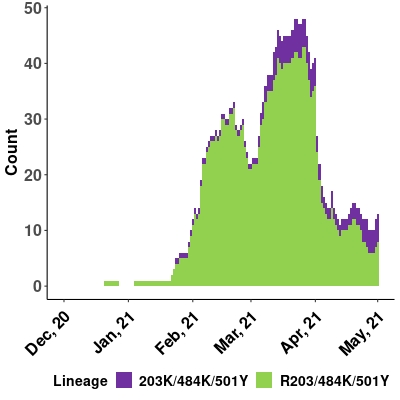

### Switzerland.jpeg

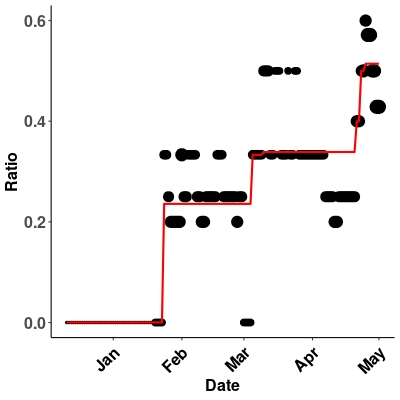

### Switzerland.jpeg

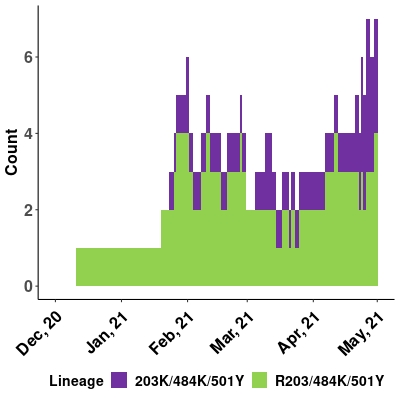

### Thailand.jpeg

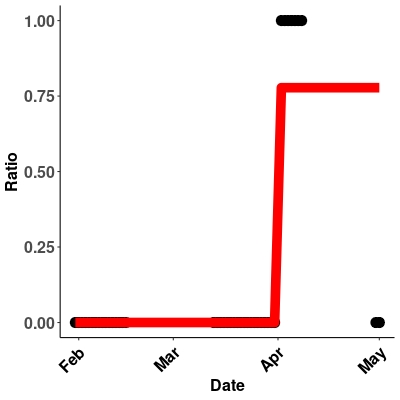

### Thailand.jpeg

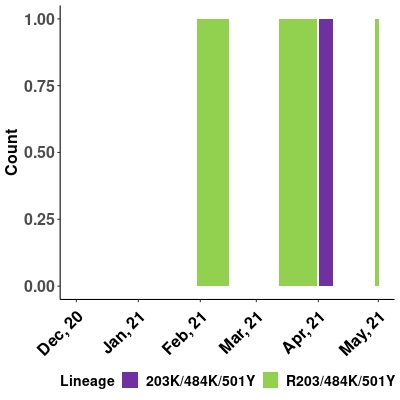

### Turkey.jpeg

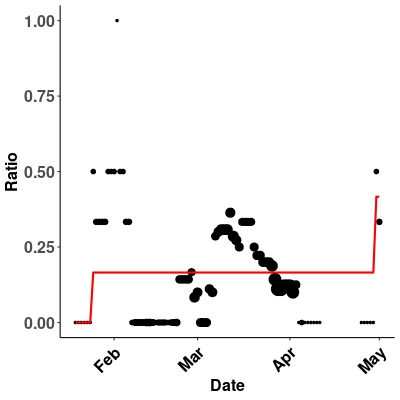

### Turkey.jpeg

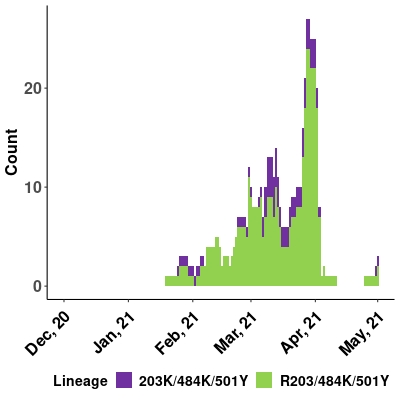

### United_Kingdom.jpeg

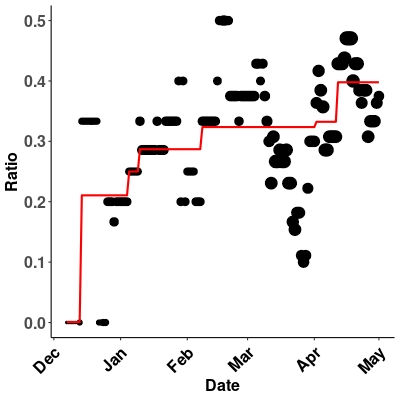

### United_Kingdom.jpeg

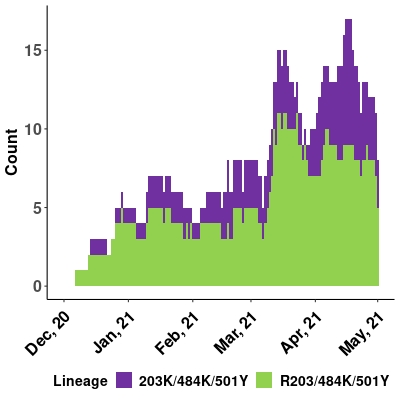

### USA.jpeg

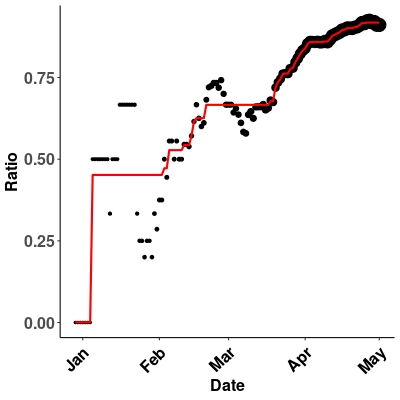

### USA.jpeg

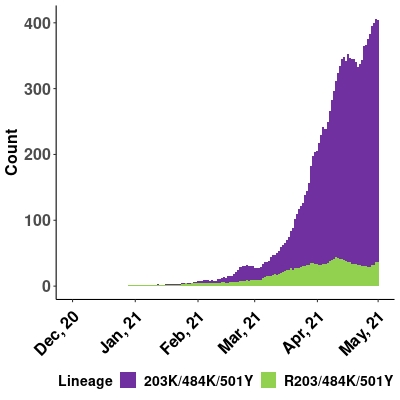
